# EHBP1 and EHD2 regulate Dll4 caveolin-mediated endocytosis during blood vessel development

**DOI:** 10.1101/2020.05.19.104547

**Authors:** Amelia M Webb, Caitlin R Francis, Jayson M Webb, Hayle Kincross, Keanna M Lundy, Rachael Judson, Dawn Westhoff, Stryder M Meadows, Erich J Kushner

## Abstract

Despite the absolute requirement of Delta/Notch signaling to activate lateral inhibition during early blood vessel development, many mechanisms remain unclear. Here, we identify EHD2 and EHBP1 as novel regulators of Notch activation in endothelial cells through controlling endocytosis of Delta-like ligand 4 (Dll4). Knockout of EHBP1 and EHD2 in zebrafish produced a significant increase in ectopic sprouts in zebrafish intersomitic vessels during development and a reduction in downstream Notch signaling. *In vitro*, EHBP1 and EHD2 localized to plasma membrane-bound Dll4 and actin independently of clathrin. Disruption of caveolin endocytosis resulted in EHBP1 and EHD2 failing to organize around Dll4 as well as loss of Dll4 internalization in endothelial cells. Overall, we demonstrate that EHBP1 and EHD2 regulate Dll4 endocytosis by anchoring caveolar endocytic pits to the actin cytoskeleton.

## INTRODUCTION

Angiogenesis is the process in which a new blood vessel develops from an existing vascular bed. During this process, in response to a chemotactic gradient, a single endothelial cell (EC) in a group must identify itself as a tip cell [1]. This tip cell will then lead the charge up the growth factor gradient, pulling behind it a trail of stalk cells in the growing vascular sprout [2]. Tip and stalk cells each have distinct morphological and functional identities: the tip defined by its spiny, branching filopodia reaching forward as the tip migrates; the stalk defined by its smoothened appearance and suppressive stability [3, 4]. For proper angiogenic growth to proceed, the maintenance of tip/stalk cell specification is paramount. Central to tip/stalk cell specification is the Notch signaling pathway. Notch is a transmembrane protein composed of an extracellular domain (NECD) and an intracellular domain (NICD). ECs with high Notch activation will adopt a stalk cell identity, whereas an EC deficient in Notch or Notch signaling will adopt a tip cell identity [5, 6].

Delta like proteins are transmembrane Notch ligands. The NECD of a Notch presenting cell will bind Delta on an adjacent cell. This Delta/Notch binding elicits two consecutive cleavage events. Obscured within two domains (LNR and HD) of Notch is a cleavage site termed S2. When exposed, the S2 cleavage site is cleaved by a disintegrin and metalloprotease (ADAM) complex leaving the NECD attached to Delta [7]. This cleavage event precedes the second cleavage by γ-secretase at the S3 cleavage site to release the NICD. Once freed, the NICD translocates to the nucleus, binding the transcription factor CSL to upregulate downstream genes that promote lateral inhibition [8–10].

One proposed mechanism for this activation of Notch by Delta is the application of a mechanical force generated by Delta/NECD transcytosis (i.e. endocytosis of both Delta and NECD) to expose the S2 domain. This pulling force has been shown to be on the order of 19 pN per single bond [11] and is necessary to force apart the LNR/HD interaction, thus exposing the S2 site for cleavage and subsequent S3 cleavage. To date, studies on the endocytic mechanisms that underly Delta/Notch transcytosis to date have only focused on Delta-like ligand 1 (Dll1) in non-endothelial tissue [11–13]. Despite the absolute requirement of Delta-like ligand 4 (Dll4) in Notch signaling, the mechanisms of Dll4 transcytosis remain unknown.

In mammals, 4 Epsin15 homology domain (EHD) EHD1-4 are each involved in endocytic processes, although EHD2 stands alone from this group in being the only with a solved crystal structure and the only to interact with caveolae [14]. EHD2 multimerizes through an interaction between the G-domain and the EH domain [14, 15]. The multimerization of EHD2 allows it to form a circular ring around an endocytic vesicle to mediate pit stability [14]. This complex localizes to caveolae, assisting in caveolin-mediated endocytosis through stabilization of the caveolae pit structure formed by hairpin shaped proteins in the membrane [16, 17]. EHD2’s binding partner, EHBP1, is much less well characterized. The NPF motifs of EHBP1 bind the EH domain of EHD2, while the CH domain of EHBP1 tethers the EHD2/EHBP1 complex to the cytoskeleton. EHBP1 and EHD2 regulate clathrin-mediated endocytosis of Transferrin and Glut4 receptors [18, 19]. Although, how EHBP1 and EHD2 function in ECs remains uncharacterized.

In this article, we identify EHD2 and EHBP1 as novel regulators of Notch activation in ECs through controlling endocytosis of Dll4. Knockout of EHBP1 and EHD2 in zebrafish produced a significant increase in ectopic sprouts in zebrafish intersomitic vessels (ISVs) during development and a reduction in downstream Notch signaling. *In vitro*, EHBP1 and EHD2 localized to plasma membrane-bound Dll4 and actin independently of clathrin. Disruption of caveolin endocytosis resulted in EHBP1 and EHD2 failing to organize around Dll4 as well as loss of Dll4 internalization in ECs. Overall, we demonstrate that EHBP1 and EHD2 regulate Dll4 endocytosis by anchoring caveolar endocytic pits to the actin cytoskeleton.

## RESULTS

### EHBP1 and EHD2 affect blood vessel development

To investigate whether the EHBP1 and EHD2 complex play a role in angiogenesis, we first characterized how the individual loss of EHBP1 or EHD2 affected intersomitic blood vessel (ISV) development in *Danio rerio* (zebrafish). This vessel bed requires tightly regulated tip/stalk cell specification and demonstrates stereotyped morphodynamics, making aberrations in normal blood vessel development relatively obvious [20]. Due to a gene duplication event in teleosts, EHD2 has two paralogs in zebrafish: EHD2a and EHD2b. We targeted each paralog individually, as well as in combination, using a 4-guide CRISPR knockout (KO) approach [21] to create F0 KOs in the EHBP1 and EHD2a/b loci. For each KO, we evaluated one of the four CRISPR cut sites for indel formation. Sequencing revealed a 100% of the putative target sites contained substantial indels in 3 random samples from each condition (Figure 1A-C). We were confident coding of each gene was significantly disrupted given the efficacy of indel formation and each gene was targeted in four separate loci. Quantification of the proportion of fish with ectopic ISVs using a vascular reporter line tg(kdrl:eGFP) [22] in each condition revealed a significant increase in EHBP1 (18.15%) and EHD2a/b knockout fish (21.76%) compared to a scrambled single-guide RNA control (Figure 1D-G) at 48 hours post fertilization (hpf). These results suggest that EHBP1 and EHD2a/b are necessary for normal sprouting behaviors.

**Figure 1.**
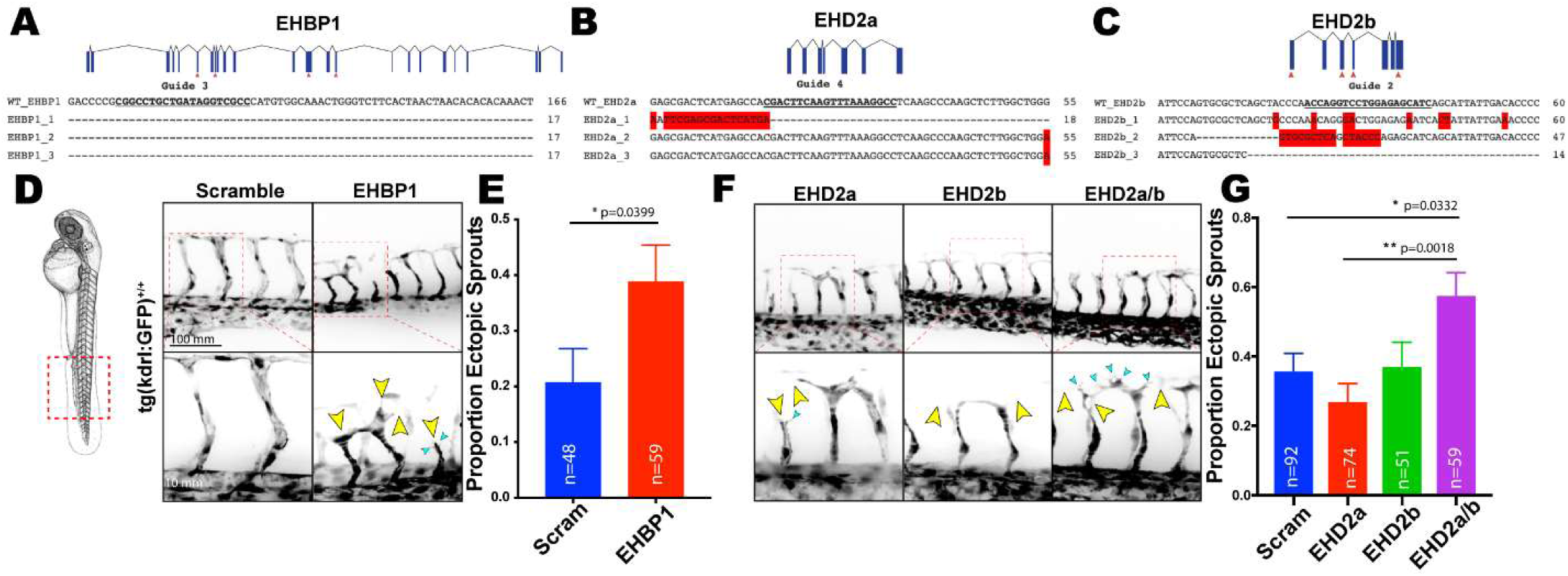
EHBP1 and EHD2 promote ectopic sprouting in zebrafish blood vessels. (A-C) Sequence alignment of NCBI sequence of EHBP1 (C), EHD2a (F), or EHD2b (G) in comparison to 4-guide KO injected fish (n=3). (D,F) Red dashed box denotes area of imaging. 4-guide knockout (KO) fish were imaged on tg(kdrl:GFP)^+/+^ background at 48 hpf. Yellow arrows denote abnormal vascular growth. Teal arrows point to spiny projections. (E,G) Quantification of the proportion of 4-guide KO fish with ectopic ISVs. Ectopic was defined as a vessel projection emerging from the ISV distinct from the central stalk. Error bars represent SEM. n= number of fish quantified.

Focusing on EHBP1, as this protein is less well characterized, we sought to determine the endogenous expression of EHBP1. To circumvent the lack of commercial antibodies detecting EHBP1 in zebrafish, we fused a FLAG epitope to the N-terminal domain of endogenous EHBP1 using CRISPR/Cas9 targeting [23] (Figure 2A). Staining for FLAG-EHBP1 endogenous expression revealed that EHBP1 strongly localized to neuronal projections (Figure 2B). To determine if ECs express EHBP1, we dissociated embryos expressing both FLAG-EHBP1 and a GFP-endothelial reporter at 48hpf and plated individual ECs on a dish. We found there was indeed EHBP1 expression in ECs at this time point; although in low abundance (Figure 2C). Although we could not detect EHD2a by *in situ* hybridization, EHD2b showed a similar expression pattern to EHBP1 along neuronal projections [24] (Figure S2A).

**Figure 2.**
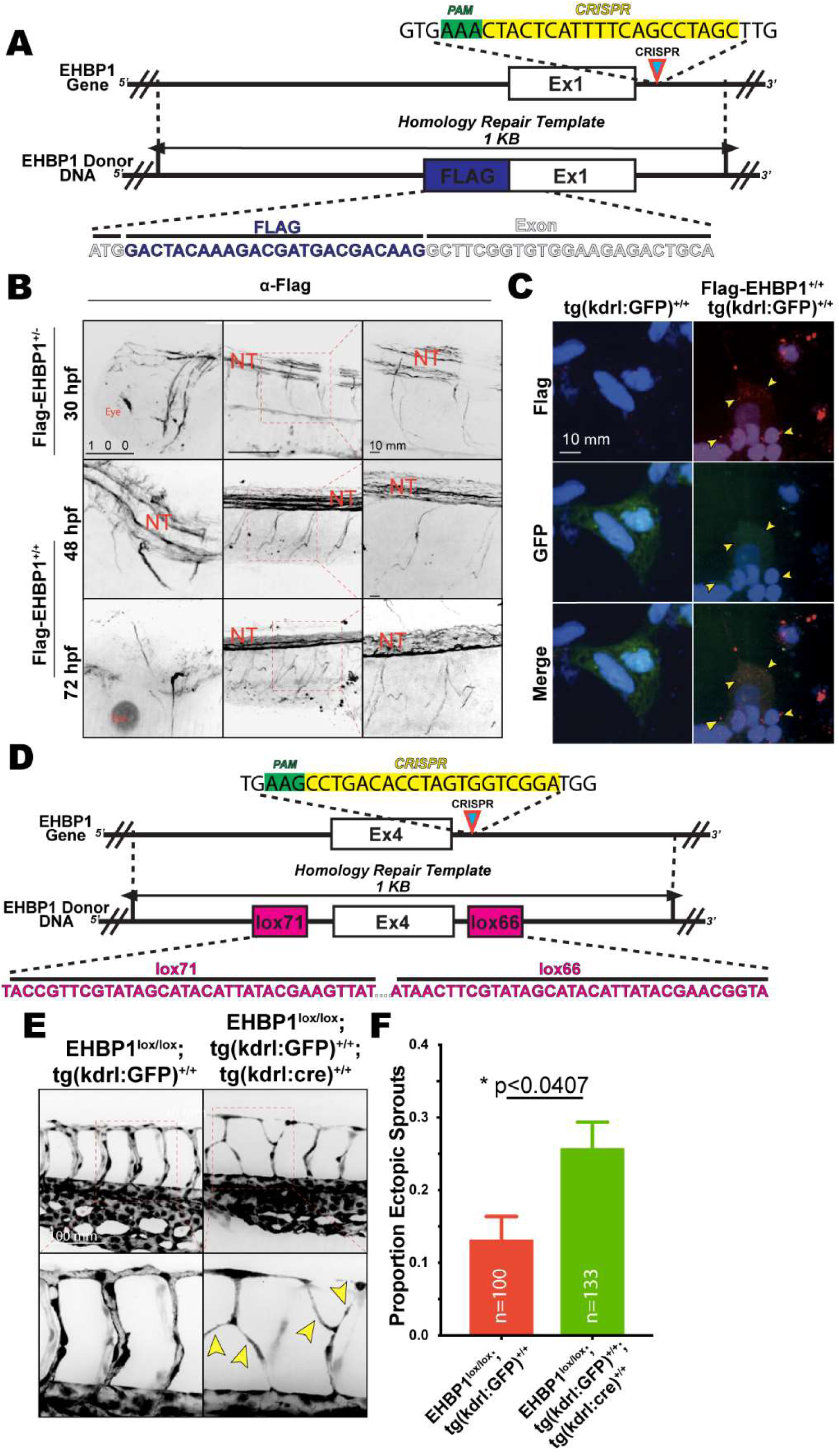
EHBP1 is expressed in the vasculature and is required for proper sprouting. (A) Targeting strategy to introduce N-terminal FLAG epitope to EHBP1 exon 1. (B) Staining against FLAG epitope in various stages of development highlighting expression in the neural tube (NT) in indicated genotypes. (C) Ex vivo validation of expression of FLAG-EHBP1 in endothelial cells. Yellow arrows denote areas of FLAG positivity. (C) Targeting strategy to introduce lox71 and lox66 sites. (E) Representative images of ISVs in indicated genotypes. Yellow arrows denote abnormal vascular growth. (F) Quantification of proportion of fish with ectopic ISVs. Error bars represent SEM. n= number of fish quantified.

Given the potentially confounding effects of performing a global KO as well as the relatively low abundance of EHBP1 in ECs, we created a floxed EHBP1 transgenic line via CRISPR/cas9- mediated recombination to test the role of this protein in angiogenesis. To do so, we flanked Exon 4 with Lox71 and Lox66 sites (Figure 2D, S1B) and crossed this line to a vascular reporter also expressing Cre recombinase [25]. In line with our previous observations, endothelial-specific, homozygous ablation of EHBP1 also resulted in a significant increase in the proportion of fish with ectopic ISVs (Figure 2E,F). These results suggest that EHBP1 and EHD2a/b mediate endothelial sprouting behaviors and are required for normal blood vessel development.

### EHBP1 and EHD2 modulate Notch signaling in zebrafish

We next tested if vascular abnormalities in EHBP1 and EHD2a/b KO lines were related to Notch activity, as loss of Notch signaling promotes hypersprouting both in developing zebrafish and mouse blood vessels [5, 26–31]. Treatment with the small molecule Notch inhibitor LY-411575 phenocopied the increase in ectopic sprouting observed in EHBP1 and EHD2 KOs (Figure 3A, B), suggesting that KO of EHBP1 or EHD2 may be detrimental to Notch signaling. We also observed an elevated number of filopodial projections characteristic of tip cell identity [3] in the LY-411575 treated fish (Figure 3A, C). We repeated the 4-guide CRISPR KO injections for EHBP1 or EHD2a/b and quantified the number of filopodia per ISV in an actin-labels EC reporter (tg(kdrl:LifeAct-eGFP) [32] line as performed above. KO of EHBP1, EHD2a, EHD2b, and EHD2a/b did not result in a significant increase in filopodia compared to a scramble control (Figure 3D, E), likely due a less severe Notch deactivation compared with LY-411575. To further explore Notch activation, we monitored expression of Hey2, a downstream Notch target, across groups in reference to a GAPDH control. We observed significantly diminished expression of Hey2 in EHBP1, EHD2b, and EHD2a/b KO lines in comparison with the scramble control (Figure 3F). The lack of an effect with EHD2a KO is likely due to the lack (or weak) mRNA expression of EHD2a at this time point. Overall, these results support a Notch loss-of-function phenotype in the absence of EHBP1 and EHD2b.

**Figure 3.**
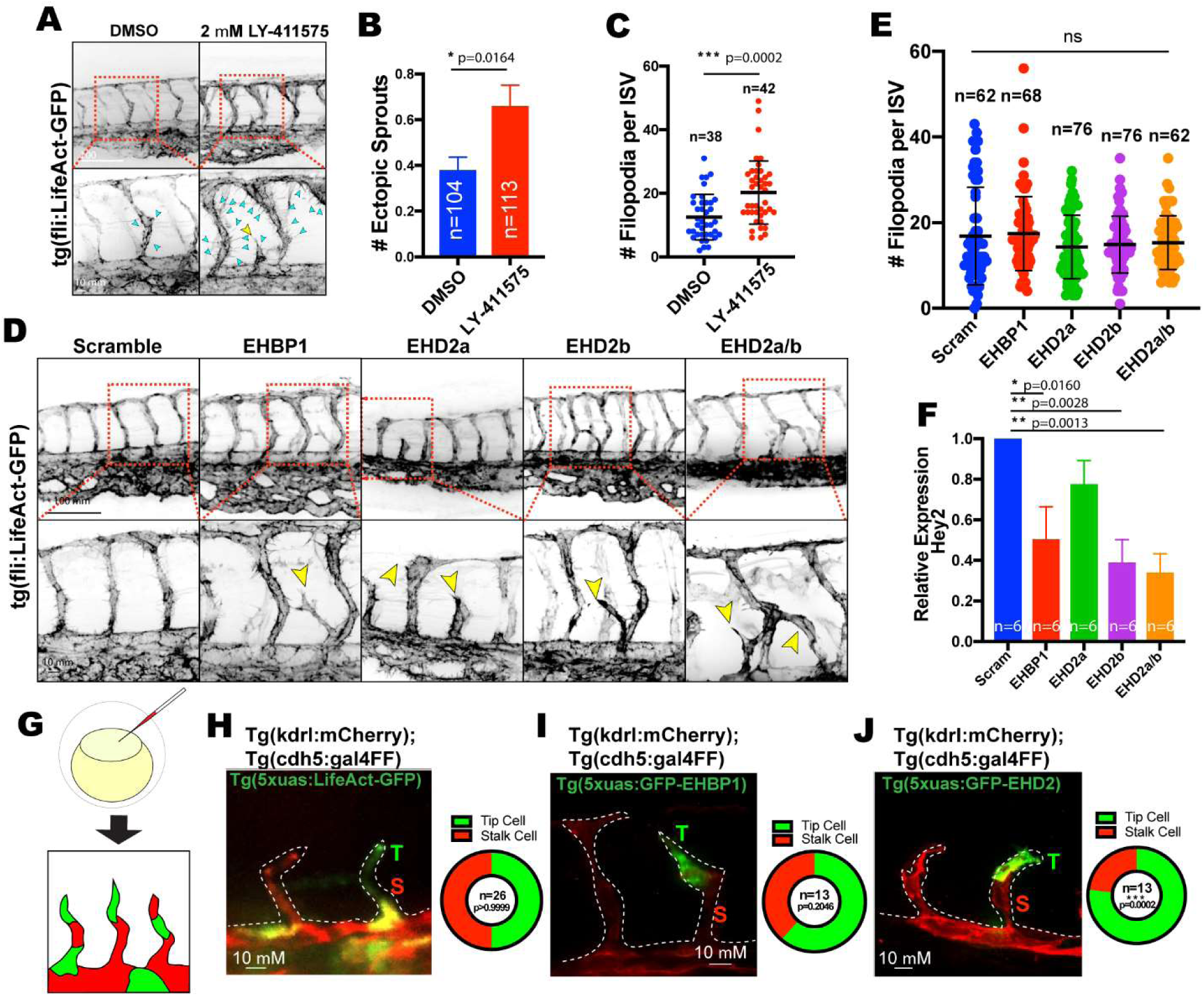
EHBP1 and EHD2 knockouts phenocopy Notch loss of function. (A) ISVs of fish treated with either DMSO or 2 µM LY-411575 on tg(fli:LifeAct)^+/+^ background. Yellow arrows point to ectopic sprouts. Teal arrows point to filopodial projections. (B) Number of ectopic ISVs per fish between indicated groups. (C) Number of ISV filopodial extensions between indicated groups. (D) Representative images of 4-guide KO of EHBP1 and EHD2 on tg(fli:LifeAct-GFP)^+/+^ background. Yellow arrows point to ectopic sprouts. (E) Number of ISV filopodial extensions among indicated groups. (F) Relative expression of Hey2 transcript in 48hpf 4-guide KO fish compared to GAPDH control across indicated groups. (G) Injected embryos will produce mosaic expression. (H-I) Representative images of LifeAct-GFP, GFP-EHBP1, and GFP-EHD2, overexpression constructs in growing ISVs at 24 hpf. Quantification of the proportion of expressing endothelial cells in either the tip (T, green) or stalk (S, red) cell position in the growing vascular sprouts shown to the right. n= number of fish, ns= non-significant. Error bars represent SEM in (B) and (F) and SD in (C) and (E).

In canonical tip/stalk cells specification, tip cells exhibit elevated Dll4 levels that, in turn, elicit repressive Notch activation in the trailing stalk cells [5, 6, 29, 30, 33, 34]. Given that KO of EHBP1 or EHD2a/b promoted a Notch loss-of-function hypersprouting phenotype, we hypothesized that these proteins may influence hierarchical tip/stalk cell positioning. To determine how EHBP1 and EHD2 function during tip/stalk cell specification, we developed GFP-tagged EHBP1 and EHD2 fusion proteins that were injected into a WT (tg(kdrl:mCherry; tg(chd5:gal4ff) [35] vascular reporter line to produce mosaic ISVs in order to visualize individual ECs in the sprout collective (Figure 3G). Zebrafish EHBP1 and EHD2a/b protein are approximately 70% identical to the Human ortholog, thus predicted to work similarly (Figure S2). We reasoned that if EHBP1 or EHD2 did not affect Notch activation there would be an equal hierarchical EC contribution in tip or stalk cell positions. Confirming this, we injected a control tg(5xUAS:LifeAct-GFP) construct and observed mosaic integration with an even 50/50 distribution between tip and stalk ECs in growing ISVs at 24hpf (Figure 3H). However, ECs expressing GFP-EHD2 (tg(5xUAS:GFP-EHD2)) demonstrated a significant bias toward the tip cell position with 76.9% of ISVs exhibiting EHD2-overexpressing ECs in the tip position (Figure 3I). Similarly, GFP-EHBP1-expressing ECs showed a 61.5% non-significant bias to the tip cell position in ISVs at 24hpf (Figure 3J). In our experience a limited number of ECs tolerated the overexpression EHBP1 *in vivo* potentially due an incompatibility between the WT and overexpressed version of EHBP1. These results support the role of EHBP1 and EHD2 in tip cell-related dynamic competition and Notch signaling.

### EHBP1, EHD2, and actin surround membranous Dll4 independent of clathrin

Given the effect of EHBP1 and EHD2 on Notch activation *in vivo*, we hypothesized that EHBP1 and EHD2 participates in Dll4 endocytosis, as Dll4-mediated signaling requires endocytosis [36]. EHBP1 and EHD2 contribute to both clathrin-dependent and independent endocytosis [14, 15, 18, 19]. To determine the endocytic route employed, we moved to an *in vitro* culture-based model using primary ECs (Human umbilical vein ECs). Here, we first focused only on the extracellular, membrane-inserted pool of Dll4 that would be available for Notch binding. To specifically label this bioactive Dll4 population we constructed a pHluorin-tagged Dll4 adenoviral vector. PHluorin is a GFP variant that fluoresces at neutral pH and is quenched when internalized into low pH acidic vesicles allowing for visualization of the extracellular, membrane bound Dll4 population [37] (Figure 4A). Co-expression of pHluorin-Dll4 and tagRFP-EHBP1 in ECs showed that EHBP1 clustered at sites of membranous Dll4 (Figure 4B). A similar pattern was observed for EHD2, where EHD2 encircled Dll4 puncta (Figure 4C). These results support the potential role of EHBP1 in Dll4 endocytosis. However, staining for endogenous clathrin revealed that clathrin was largely divorced from Dll4 sites (Figure 4B,C). Regression analysis of the proportion of clathrin and EHBP1 present at Dll4 puncta demonstrated a strong positive correlation (pearson’s coefficient 0.9049), suggesting that if clathrin was present at Dll4 puncta, so was EHBP1 (Figure 4D), indicating that EHBP1 may have a dual role in both clathrin-dependent and independent endocytic processes, or EHBP1 is present on puncta that are not actively undergoing endocytosis. Interestingly, there was no association between clathrin and EHD2 (pearson’s coefficient 0.0363) at Dll4 puncta indicating these events were mutually exclusive (Figure 4E).

**Figure 4.**
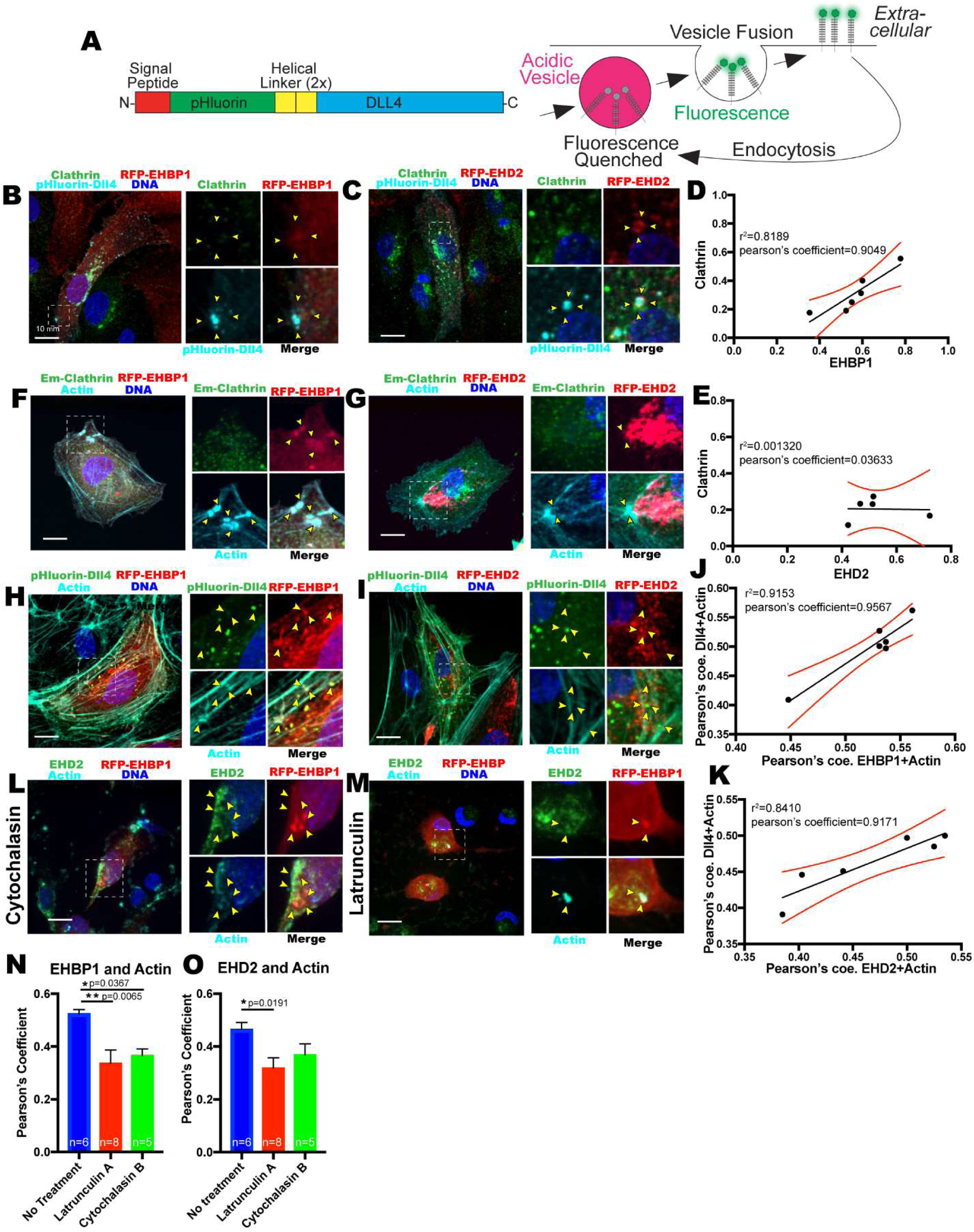
Endothelial Dll4 does not localize with EHBP1 and EHD2 on clathrin endocytic pits. (A) Schematic of pH-dependent function of GFP variant pHluorin tag. PHluorin fluoresces on the membrane at neutral pH, but is quenched when internalized into acidic endosomes. (B) EHBP1 localizes around sites of pHluorin-Dll4 independent of clathrin. (C) EHD2 localizes around sites of pHluorin-Dll4 independent of clathrin. (D) Proportion of coincidence of clathrin (y-axis) and EHBP1 (x-axis) around Dll4 puncta. (E) Proportion of coincidence of clathrin (y-axis) and EHD2 (x-axis) around Dll4 puncta. (F,G) EHBP1 and EHD2 localized to sites of F-actin bundling independent of clathrin. (J,K) Proportion coefficient between Dll4 and Actin (y-axis) and EHBP1(x-axis, J) or EHD2(x-axis, K) and actin. (H,I) EHBP1 and EHD2 cluster around Dll4 puncta at sites of F-actin bundling. (L,M) Actin inhibitors Cytochalasin B and Latrunculin A disrupt EHBP1 and EHD2 localization. (N,O) Pearson’s coefficient of EHBP1 (N) or EHD2 (O) between indicated treatments. Error bars represent SD. Boxed are magnified section on right. Yellow arrows show areas of colocalization. What is your sample size here for this analysis? Is it n=6 (6 independent transfections)?

Both EHBP1 and EHD2 demonstrated a strong association with Dll4 at F-actin bundles as previously reported [18], but clathrin was principally absent in these areas (Figure 4F,G). In probing for EHBP1, EHD2 and Dll4 puncta (Figure 4H,I) we observed a strong association between extracellular Dll4 puncta and EHBP1 or EHD2 on actin filaments (pearson’s coefficient 0.9567 and 0.9171, respectively; Figure 4J,K), indicating a potential preference of Dll4 endocytosis over actin-rich areas. Treatment of ECs with actin inhibitors cytochalasin B or latrunculin A ablated the colocalization between EHBP1 and EHD2 (Figure 4L-O). Upon treatment, localization of EHBP1 and EHD2 became diffuse and non-specific (Figure 4L,M). At select sites where actin filaments persisted, there was a slight localization of EHBP1 and EHD2. In aggregate, these data suggest that both EHBP1 and EHD2 associate with membranous Dll4 and anchor on actin filaments.

Next, we explored the notion that EHBP1 and EHD2 might coordinate Dll4 endocytosis through caveolin endocytosis as the association of clathrin to Dll4 sites was relatively weak. EHD2 contains a dynamin-like ATPase to drive the scission of caveolar endocytic pits from the membrane, although this role has not been linked to Dll4, EHBP1, or actin-tethering. TagRFP-EHBP1 and endogenous caveolin-1 both formed around clusters of pHluorin-Dll4 on the membrane (Figure 5A). Caveolin and EHBP1 demonstrated a stronger association with Dll4 (pearson’s coefficient 0.9262) (Figure 5B) as compared with clathrin (Figure 4D). Equally, TagRFP-EHD2 and caveolin-1 both formed clusters of rosettes around the Dll4 puncta (Figure 5C), also demonstrating a strong correlation (Figure 5D). To determine if this colocalization was indicative of caveolar endocytosis, we used the cholesterol inhibitor methyl-β-cyclodextrin (MβCD) to inhibit caveolar assembly. Caveolin-1, EHBP1 and EHD2 in ECs treated with MβCD all failed to form complexes around Dll4 and randomly dispersed in the cytoplasm (Figure 5E, F). To further confirm that EHBP1 and EHD2 use caveolae to internalize Dll4, we specifically knocked down caveolin-1 using siRNA. Loss of caveolin resulted in reduced EHD2 recruitment to Dll4 puncta (Figure 5G,H). Of note, knockdown of required clathrin components, clathrin-light chain B or AP2 (AAK1), did not disrupt EHBP1 or EHD2 localization to Dll4 (Figure 5I,J). These results suggest that EHBP1 and EHD2 control Dll4 internalization through caveolin-mediated endocytosis.

**Figure 5.**
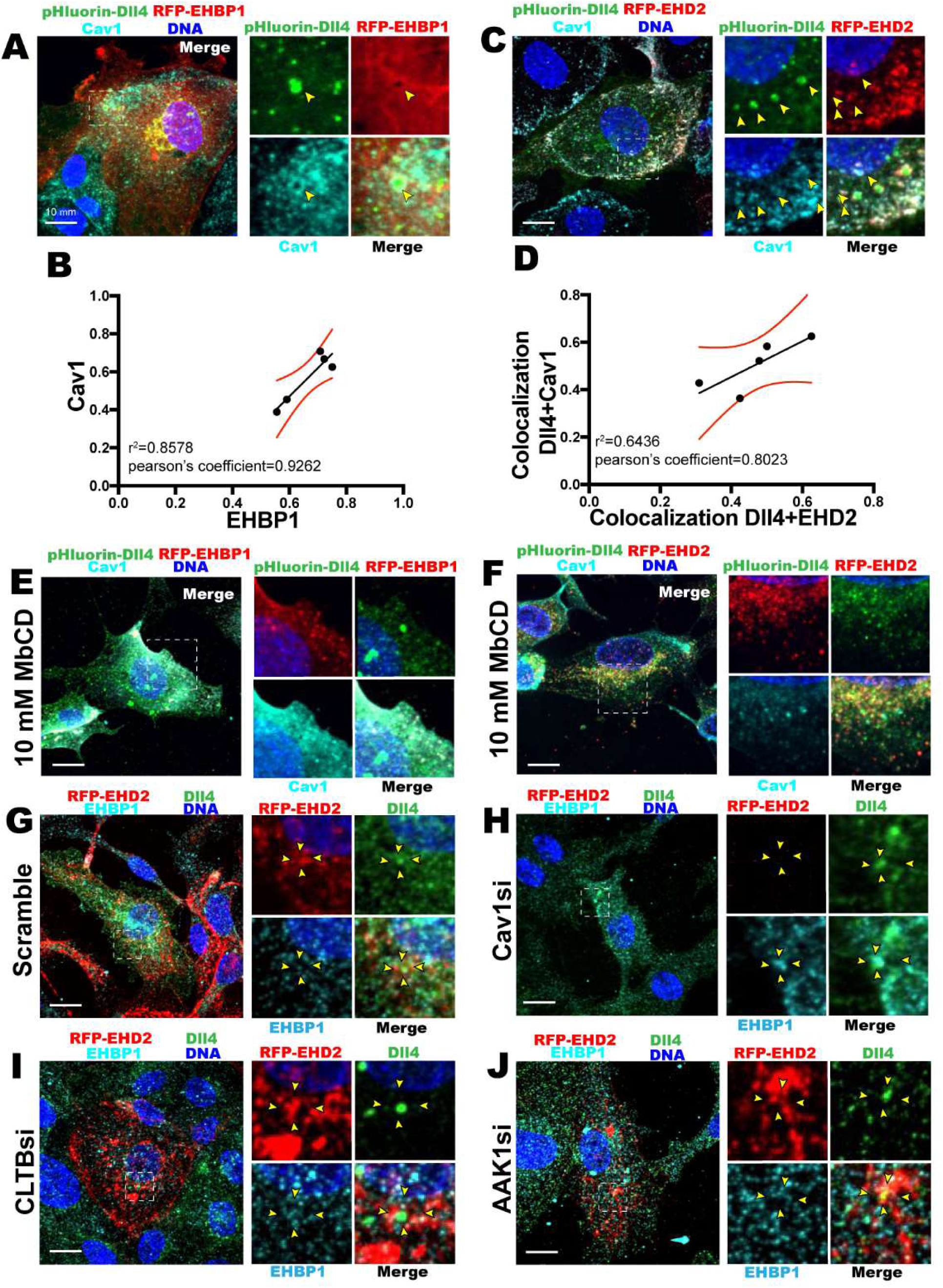
Endothelial Dll4 localizes with EHBP1 and EHD2 on caveolin endocytic pits. (A,C) EHBP1 and EHD2 localize to surface pHluorin-Dll4 and colocalize with caveolin-1. (B,D) Proportion of coincidence of caveolin (y-axis) and EHBP1 (x-axis, B) or EHD2 (x-axis, D) around Dll4 puncta. (E) EHBP1 and (F) EHD2 fail to localize to Dll4 when treated with MβCD (cholesterol inhibitor). (G-J) Expression RFP-EHD2 is reduced with Cav1si treatment. Boxed are magnified section on right. Yellow arrows show areas of colocalization. Same comment about sample size.

To validate the contribution of EHBP1, EHD2, and caveolin-1 in regulating Dll4 endocytosis in an angiogenic context, we employed a 3D sprouting assay [38] (Figure 6A). In this assay, ECs undergo collective migration making multicellular sprouts that branch and lumenize, faithfully mimicking *in vivo* processes [39–41]. EHBP1 strongly colocalized with actin in 3D sprouts consistent with reports that EHBP1 tethers actin through its CH domain (Figure 6B) [18]. This localization was disrupted in sprouts treated with EHD2 siRNA, suggesting that EHD2 plays a role in either recruiting EHBP1 to actin or in stabilizing the connection (Figure 6C). EHBP1 displayed a strong association to VE-cadherin adherens junctions in the 3D sprout (Figure 6D,E), which was also severely disrupted in EHD2 siRNA treated ECs (Figure 6F,G). Equally, EHD2 robustly localized to adherens junctions (Figure 6H,K), which were disrupted by EHBP1 siRNA knockdown (Figure 6I,L). These results support the notion that EHBP1 and EHD2 localize at junctions, which are areas of Dll4/Notch1 binding interactions and Dll4/Notch1 transcytosis.

**Figure 6.**
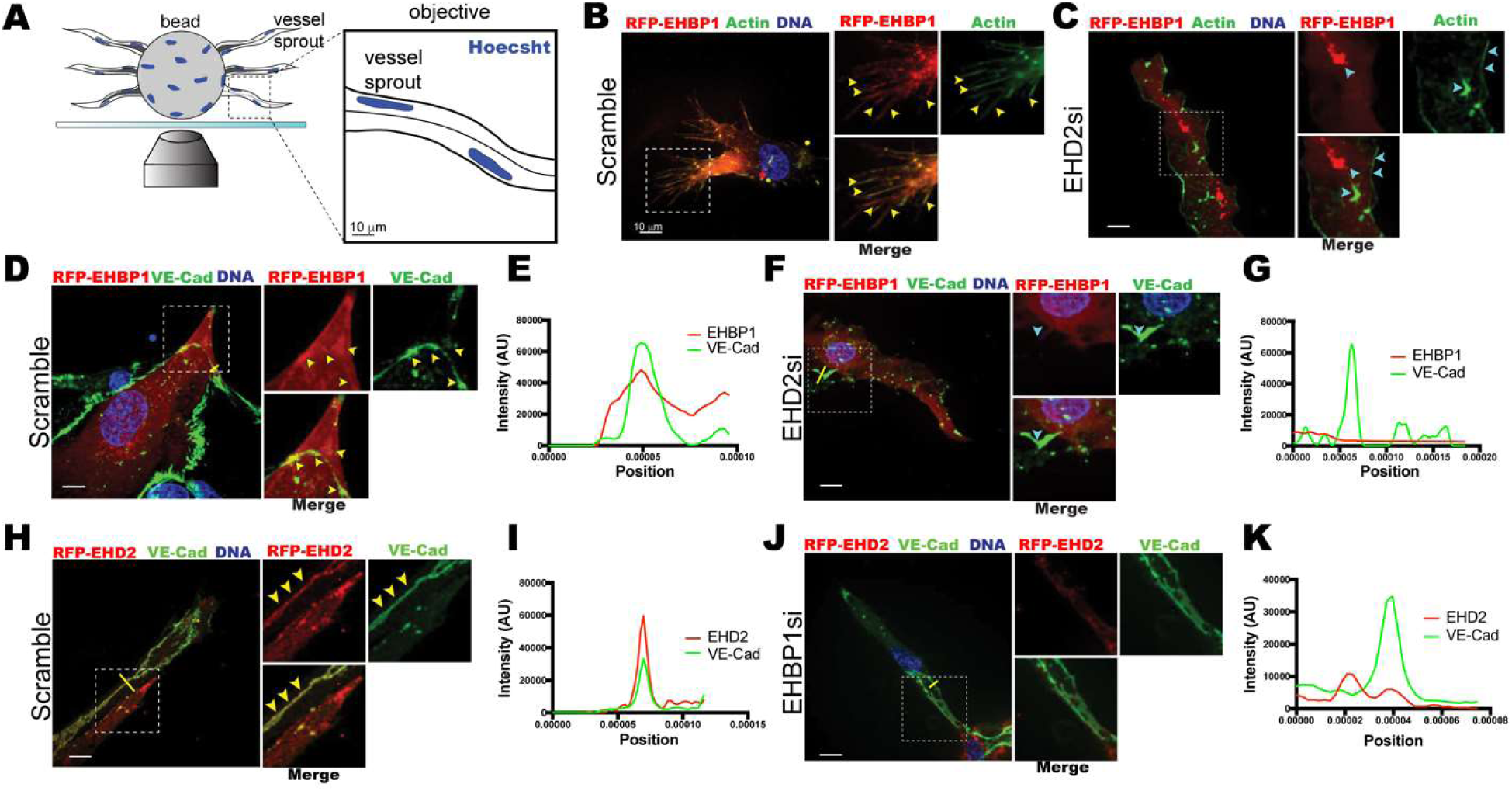
EHBP1, EHD2, and caveolin localize to adherens junctions in 3D sprouts. (A) Schematic of 3-dimensional sprout growth in fibrin bead assay (FBA). (B,C) Representative EC sprout stained for actin expressing tagRFP-EHBP1, treated with indicated siRNA. (D-G) Representative EC sprout stained for VE-cadherin (VE-cad) expressing tagRFP-EHBP1, treated with indicated siRNA. (H-L) Representative EC sprout stained for VE-cadherin (VE-cad) expressing tagRFP-EHD2, treated with indicated siRNA. Boxed are magnified section on right. Yellow arrows show areas of colocalization. Teal arrows show areas of mislocalization. What kind of cells are being used?

To confirm these interactions *in vivo* we turned to the mouse model of retinal development [42]. In E9.5 embryos and P7 retinas *in situ* hybridization staining EHD2 demonstrated elevated transcript levels in the vasculatures as well as in the developing retinal blood vessels (Figure 7A,B). In the retinal there was an enrichment of EHD2 at the vascular front. Staining for endogenous EHD2 in the retinal vasculature revealed a colocalization with Dll4 as well as caveolin-1 at the vascular front (Figure 7C,D). In our hands, multiple EHBP1 antibodies failed to detect native EHBP1 in human EC culture or in mouse tissues via immunofluorescence, although mRNA and protein expression was present via western blot. Additionally, we could not detect EHBP1 via *in situ* hybridization in mouse retina tissue potentially due to its low abundance, such as in zebrafish ECs. To this end, comparison of mouse brain and lung single EC sequencing data [43] indicated a 25 fold increase in EHD2 compared to EHBP1 (Figure S3). This is not totally surprising as EHD2 represents a more ubiquitous component of the caveolae structure, while EHBP1 may function as a more specialized endocytic adaptor.

**Figure 7.**
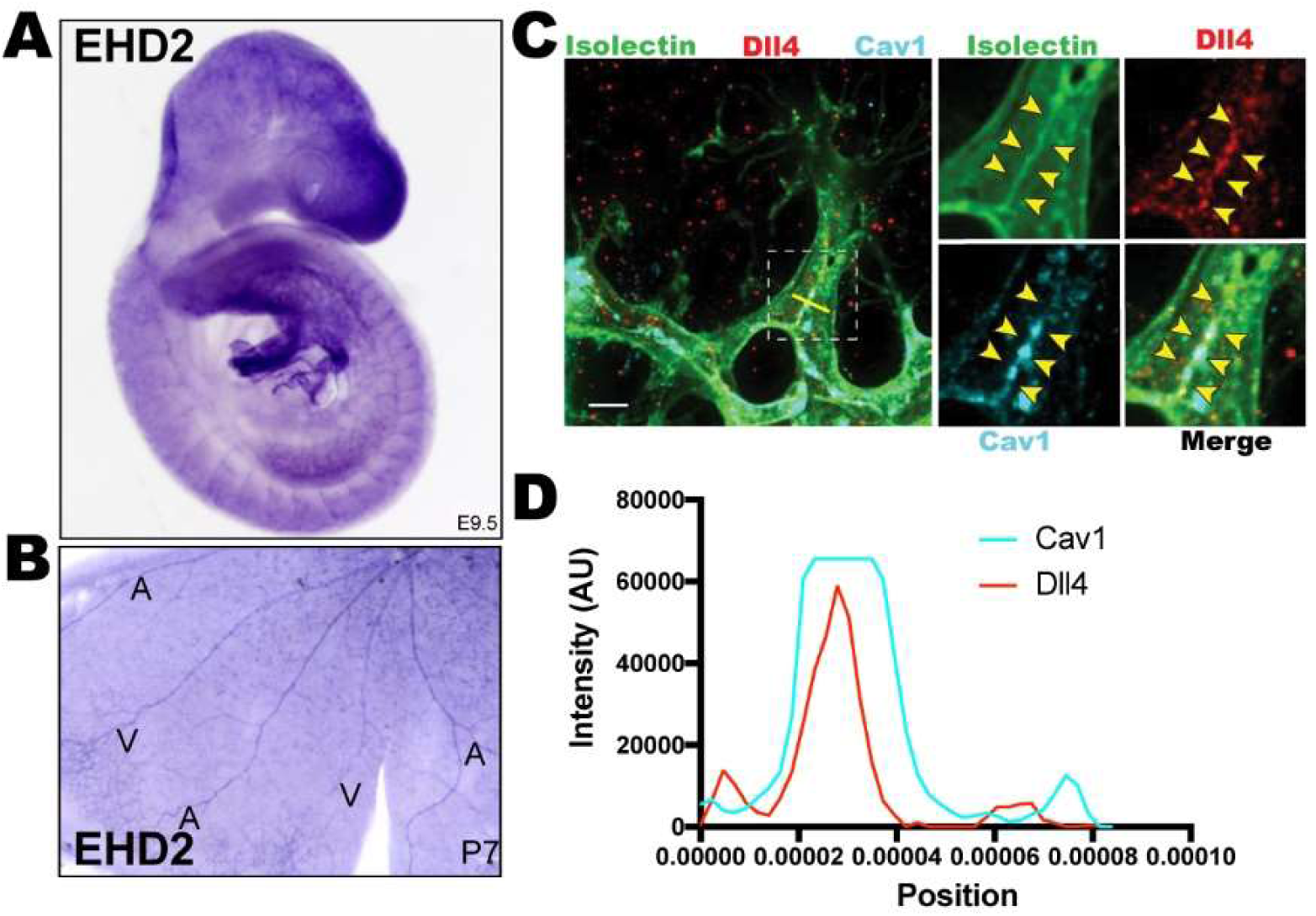
EHD2, Caveolin1, and Dll4 localize to vasculature in mice. (A-B) In-Situ Hybridization of EHD2 in whole mouse embryo (E9.5) and mouse retina (P7) (C) Staining of Isolectin, Dll4, and Cav1 in P7 mouse retinas. (D) Line scan of selected area in panel C. Boxed are magnified section on right. Yellow arrows show areas of colocalization.

### EHBP1 and EHD2 are required for Dll4/Notch transcytosis

With evidence of EHBP1 and EHD2 working together to anchor Dll4-containing caveolar pits to the cytoskeleton, we next sought to characterize Dll4 endocytic defects resulting from loss of EHBP1 or EHD2 (Figure 8A,B). To do so, we relied on a Dll4 antibody covalently linked to pHrodo, a pH sensitive dye that increases fluorescent intensity with increasing endosomal pH [44] to label extracellular Dll4, and then monitor live Dll4 endocytosis (Figure 8C, Figure S4). Pulse-chased with the pHrodo-Dll4 in control ECs, there was a sharp peak in fluorescent intensity at the 10-minute time point, indicating an increase in Dll4 endocytosis (Figure 8D). A pHrodo-labelled IgG control was added to monitor non-specific uptake. This internalization event was significantly reduced in EHBP1 siRNA, EHD2 siRNA, and combination EHBP1/EHD2 siRNA treated groups (Figure 8D). Equally, internalization of pHrodo-Dll4 in ECs treated with the pan-endocytosis inhibitor Dynasore or caveolae-specific cholesterol inhibitor MβCD also significantly reduced Dll4 internalization as compared with DMSO control (Figure 8E). These results indicate that internalization of Dll4 requires EHBP1 and EHD2 and is dynamin and caveolin dependent.

**Figure 8.**
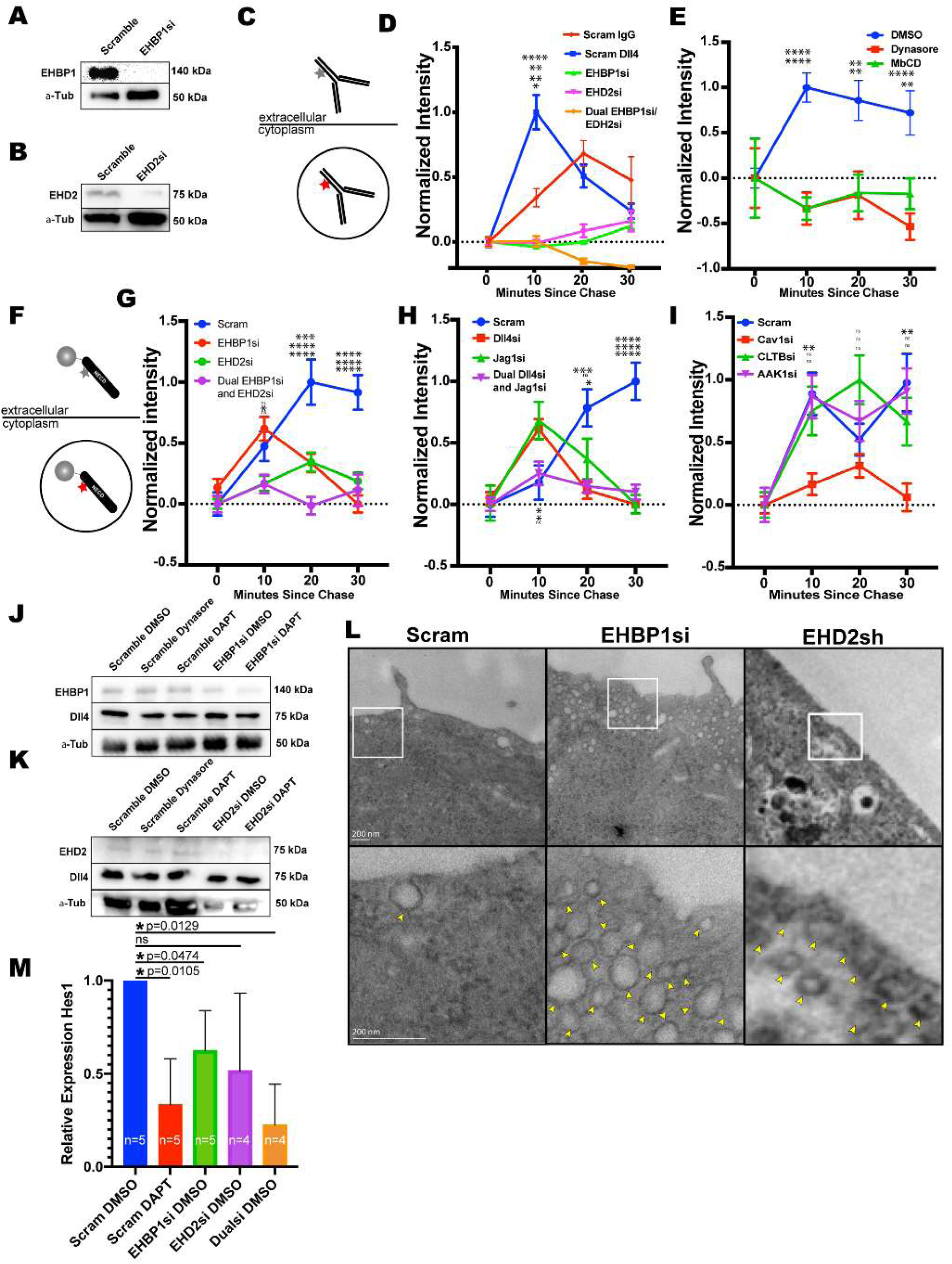
Loss of EHBP1 and EHD2 blunts Dll4 endocytosis. (A-B) Expression of EHBP1 or EHD2 after siRNA treatment in comparison to Scramble control. (C) Schematic of pHrodo labeled Dll4 antibody. PHrodo gains fluorescent intensity with increasing endosomal pH, thus used as a metric of endocytosis. (D,E) Relative internalization of pHrodo-Dll4 pulse-chase over time between indicated siRNA groups. IgG was used a non-specific internalization control. Dynasore is pan-endocytosis inhibitor. MβCD is a cholesterol specific inhibitor that disrupts caveolin. (F) Schematic recombinant Notch intracellular domain (NECD) functionalized to microbead and labeled with pHrodo. (G-I) Relative internalization of NECD pulse-chase over time between indicated siRNA groups. (J,K) Western blot of Dll4 levels across indicated treatment groups. (L) Relative expression of Hes1 compared to GAPDH control in indicated siRNA treatment groups. (M) TEM analysis of caveolar structure in ECs in across indicated siRNA treatments. White inset boxes show area of magnification. Yellow arrowheads indicate endocytic vacuoles. Significance between scramble group and other groups listed in order of figure legend. *=p<0.05, **=p<0.01, ***=p<0.001, ****=p<0.0001. ns=non-significant.

To more unambiguously test the Notch/Dll4 binding interaction, we next pHrodo-labelled recombinant NECD protein. During physiological Dll4/Notch1 binding, NECD binds to the adjacent Dll4-presenting cell prior to S2 cleavage and transcytosis. To best reproduce this complex, we functionalized the pHrodo-labeled NECD to a microbead as previously reported [12] (Figure 8F). The analysis revealed a delay in the kinetics of internalization in scramble-treated ECs compared to the Dll4-antibody internalization, likely due to the presence of a bead tether (Figure 8G). Nonetheless, siRNA knockdown against EHBP1, EHD2, or EHBP1/EHD2 combination groups led to a significant impairment of NECD internalization (Figure 8G). We observed the same internalization defect in Dll4 siRNA, Jag1 siRNA, and dual Dll4 and Jag1 siRNA treated ECs, suggesting NECD is specific binding to these Notch ligands (Figure 8H). Lastly, to again confirm Dll4 uptake depends on caveolin, we knocked down both caveolin and clathrin-related endocytic components. Knockdown of caveolin-1 significantly reduced NECD internalization, while knockdown of clathrin-light chain or AP2 did not affect Dll4 internalization compared with control (Fig. 8I). Overall, these data indicate that EHBP1 and EHD2 are required for caveolin-mediated Dll4 endocytosis.

We considered that the disruptions in Dll4 endocytosis may be due to reduced Dll4 bioavailability. However, we observed that EHBP1 siRNA, EHD2 siRNA, DAPT, or Dynasore treatment did not affect Dll4 levels (Figure 8J, K). Therefore, the reduced internalization of Dll4 is a direct result of loss of EHBP1 and EHD2, supporting their role as endocytic mediators of Dll4/Notch1 transcytosis. In order to visualize this endocytic impairment in the absence of EHBP1 and EHD2 we employed TEM imaging. Both EHBP1 and EHD2 knockdown greatly increased the number of small endocytic vacuoles near the plasma membrane (Figure 8L), an observation consistent with previous reports investigating EHD2 [14, 45] in which caveolae are unable to be stabilized through actin anchoring and accumulate near the plasma membrane; however, this has not been shown for EHBP1. Lastly, we confirmed that similar to *in vivo* results, knockdown of EHBP1 or double EHD2/EHBP1 resulted in a significant reduction in Hes1 expression, indicating a reduction in Notch signaling (Fig. 8M).

### RNA-seq analysis of EHBP1 deficient cells

We were intrigued by the idea that EHBP1 may be a specific adaptor for caveolin-mediated endocytosis as it was less abundant, engaged only a subset of membranous Dll4 and uncharacterized in in endothelial tissue. To gain better insight into EHBP1’s role in endothelial signaling we preformed RNA-seq analysis. We evaluated KD efficiency by western blot (Figure 9A). In the EHBP1si treated group, we only identified 99 differentially regulated genes compared to the control (Table S1). A volcano plot shows the log-fold-change of gene expression between the scramble and EHBP1 siRNA treated ECs, with upregulation presented above the line and downregulation presented below the line (Figure 9B). Of these 99 differentially regulated genes, approximately 45% either contained EGF repeats, analogous to Dll4 and Notch, or were indirectly related to EGF-containing proteins signaling processes (Figure 9B, C; Table S2). Further analysis revealed that 8 of the 99 genes relates to caveolin endocytosis (Figure 9B, D) and 4 of the 99 genes function as actin remodeling genes (Figure 9B, E). This suggests that EHBP1’s role in regulating caveolar endocytosis may not be specific to Dll4, *per se*, but can accommodate a variety of cargo with an emphasis on proteins containing EGF repeats. To determine if this trend translated to an increased consequence to angiogenic signaling, we grouped the differentially regulated genes by gene ontology. 8 genes associate to either angiogenesis, blood vessel development, or response to hypoxia (Figure 9F). Another 11 genes belong to the trafficking ontologies SNARE binding, G protein-coupled receptor binding, exocyst, and MAPK cascade (Figure 9G). Beyond these, all differentially regulated gene ontologies represented in the bar plot show a dramatic impact on cellular processes as a whole (Figure 9H). Overall, EHBP1 shows an affinity for endocytic processes related to EGF-repeat containing proteins, such as Dll4.

**Figure 9.**
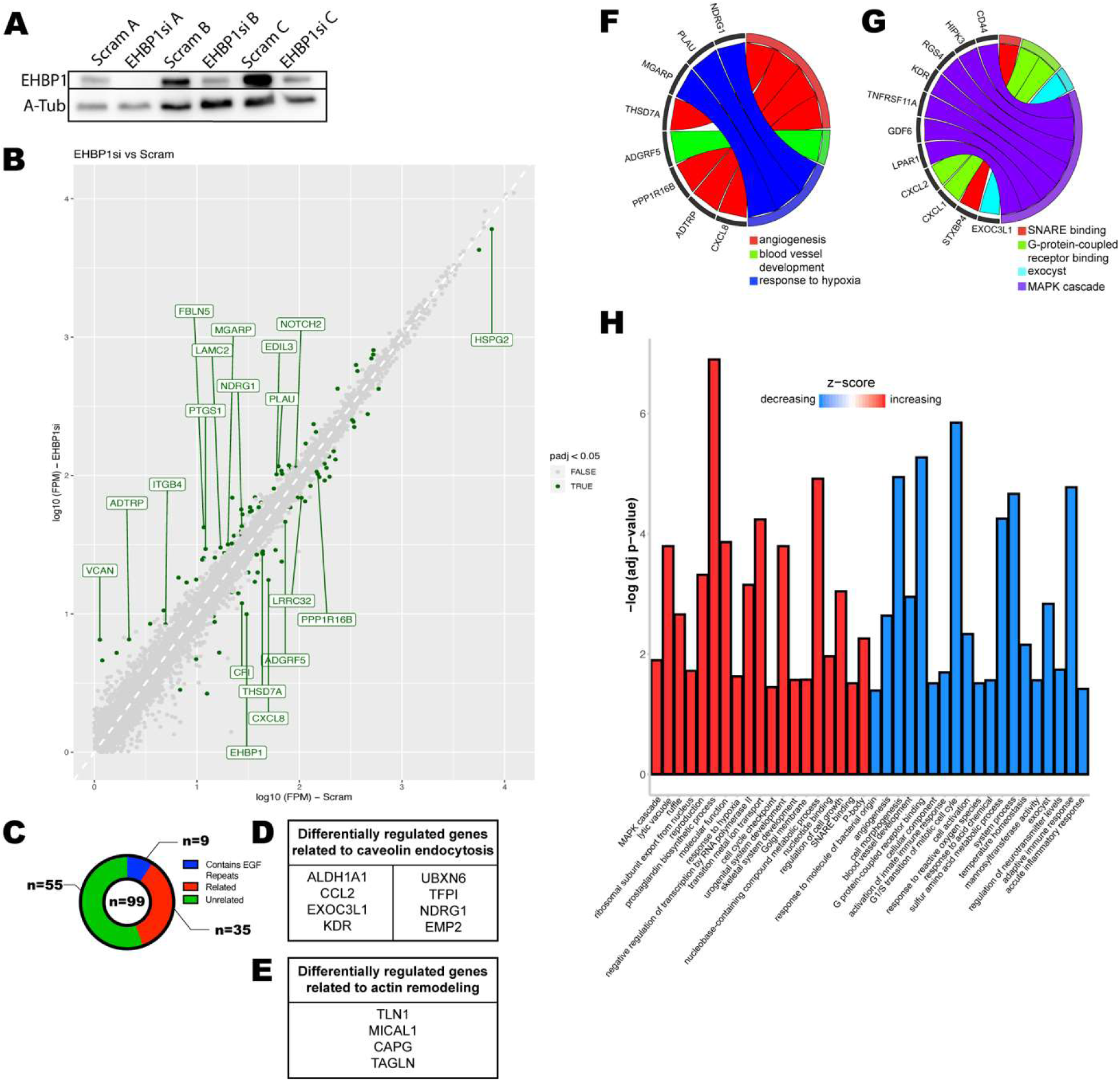
RNA-seq Analysis of EHBP1si treated cells. (A) Confirmation of knockdown efficiency of EHBP1 siRNA replicates. (B) Volcano plot distribution of all differentially regulated genes in EHBP1si condition compared to control. Dots above the line represent upregulation; dots below the line represent downregulation. Dark green dots represent significantly differentially regulated genes. Highlighted points are relevant to angiogenic signaling. (C) Number of significantly different transcripts that contain EGF repeat motifs, are (D) directly related to caveolin-mediated endocytosis or (E) directly related to actin remodeling. (F) Significantly differentially regulated genes related to angiogenesis. (G) Significantly differentially regulated to endo-or exocytosis. (H) All gene ontology groupings of significantly differentially regulated genes. Bars shown in red were upregulated as a group. Bars shown in blue were downregulated as a group.

## DISCUSSION

Although Notch signaling is critical for blood vessel development, endocytic mechanisms that regulate both Dll4 cell surface expression and Notch receptor activation remain elusive. We report that EHBP1 and EHD2 work in concert to regulate Notch signaling through the transcytosis of the Dll4/Notch1 complex by caveolar-mediated endocytosis. Importantly, this is the first characterization of Dll4 as being internalized by caveolin-mediated endocytosis. We also demonstrate that EHBP1 and EHD2 work cooperatively to regulate Dll4 endocytosis through tethering to the actin cytoskeleton. In a broader context, our results demonstrate novel endocytic pathway that directly impacts Dll4/Notch signaling which is required for proper blood vessel development.

We propose a model wherein EHD2 oligomerizes around the neck of an endocytic vesicle internalizing Dll4 (Figure 10). The EH domain of EHD2 also binds to the NPF motifs of EHBP1; EHBP1 is tethered to the actin cytoskeleton (Figure 10). In the absence of EHBP1 or EHD2, the caveolar pit will be incapable to tether to the actin cytoskeleton and will lose structure provided by the underlying actin network. This lack of pit anchoring through loss of EHBP1 or EHD2 will preclude the force necessary to initiate Dll4/NECD pulling and subsequent S2 cleavage and transcytosis. Without NECD transcytosis, γ-secretase is unable to cleave at the S3 intracellular site, and thus NICD cannot exert its function as a transcription factor, effectively blocking Notch signaling. Overall, both EHBP1 and EHD2 provide a key function in controlling Dll4-caveolar dynamics during Notch binding interactions.

**Figure 10.**
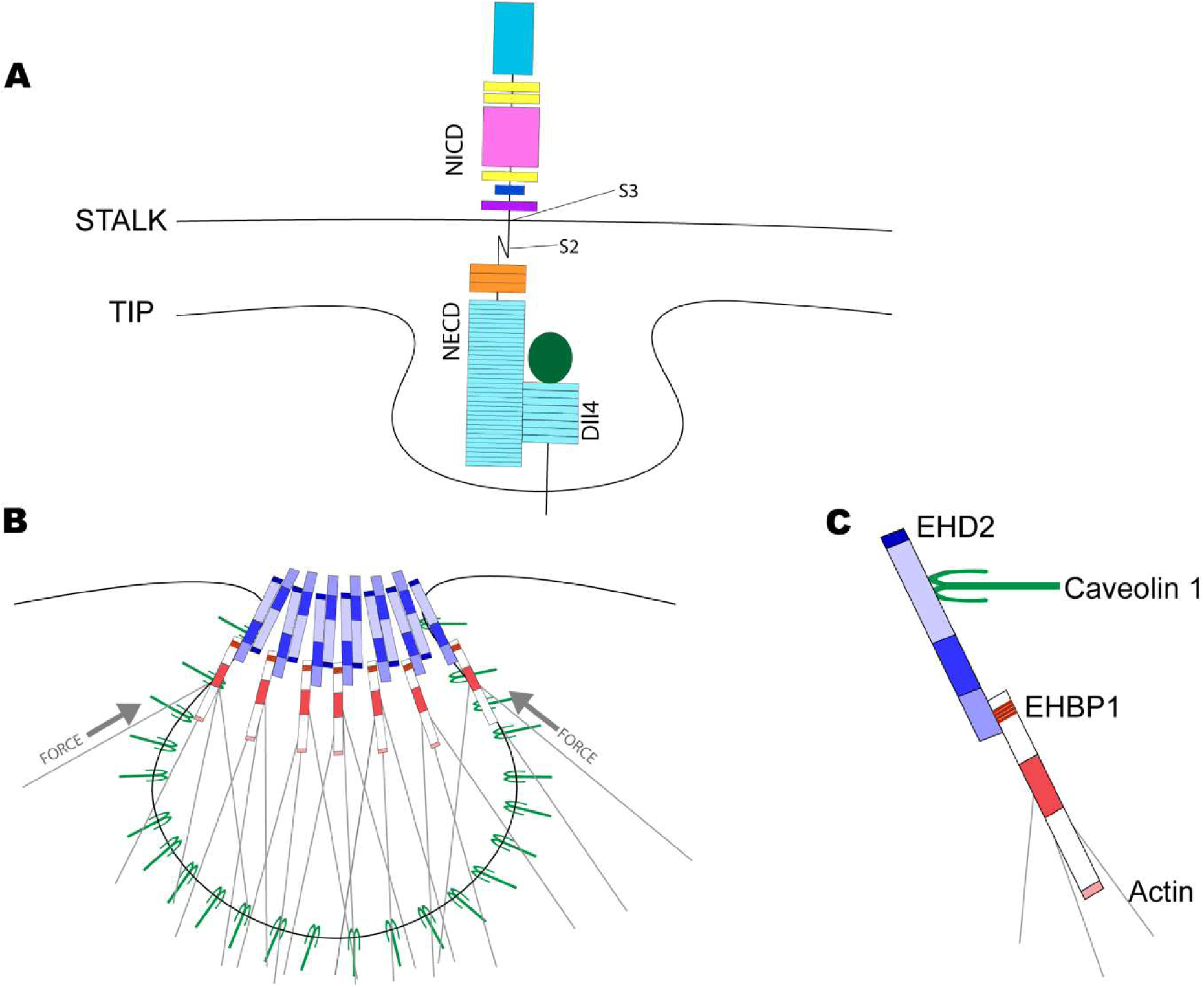
Model of EHBP1 and EHD2 mediated Dll4 internalization. (A) Dll4 binds NECD in the budding caveolin pit. (B,C) EHD2 binds caveolin and oligomerizes around the neck of the endocytic pit. EHBP1 binds to EHD2 and tethers to the actin cytoskeleton through the CH domain. This may generate the force necessary to expose the S2 cleavage site of Notch.

Dll4 is unquestionably vital for Notch signaling and blood vessel morphogenesis. In non-endothelial cell types, Dll1 endocytosis generates the mechanical pulling force on the NECD to expose the S2 domain for cleavage [11]. After S2 cleavage, the Dll1/NECD complex internalizes and, presumably, undergoes subsequent lysosomal degradation. In the adjacent Notch presenting cell, following S2 cleavage and NECD release, the S3 cleavage by γ-secretase and NICD release can proceed [7]. Others have reported that the Dll1 pulling force is derived from clathrin-dependent endocytosis [11]; however, our results suggest a different pathway. In our investigation, we equally explored the idea that EHBP1 and EHD2 use clathrin-dependent or independent programs to aid in Dll4 internalization. To our surprise, we did not observe any significant Dll4, EHPB1 or EHD2 colocalization with clathrin in ECs, indicating this was not the operative pathway for Dll4 endocytosis. Moreover, ablation of clathrin itself or related protein AP2 did not affect Dll4 endocytosis. In lieu of these results, we determined that Dll4 strongly relied on caveolar endocytosis for NECD transcytosis and subsequent Notch activation. To our knowledge, this is the first report indicating an association between Dll4 and caveolin-mediated endocytosis and it contrasts from reports investigating Dll1 [11, 12, 46, 47]. The reasons for this disparity could be due to both receptor-type and/or tissue source differences. For instance, global Dll1 deletion in mice does not affect viability, while global loss of Dll4 is embryonic lethal.

Caveolae have demonstrated a complex role in regulating both endocytosis events and maintaining tissue integrity. Caveolae are 50-80nm flask shaped invaginations of the plasma membrane shaped through integral membrane protein caveolins and their associated proteins [48, 49]. In addition to endocytosis, caveolins more recently have been shown to be membrane reservoir that can safeguard against mechanical stress [50]. For instance, caveolin 1,3 are essential for notochord integrity in developing zebrafish [50]. Similarly, caveolins protect against membrane rupture in ECs and skeletal muscle [51, 52]. Interestingly, global deletion of caveolin1 is not embryonic lethal [53], suggesting that caveolins are not essential and/or other redundant factors are at play. In our investigation, loss of either EHD2 or EHBP1 did not affect caveolin formation on the plasma membrane, although, it did impact the number of caveolar pits adjacent to the plasma membrane. Others have shown that loss of EHD2 does not affect the number of caveolar pits formed in non-endothelial tissues, but drastically increases their dynamics by not being anchored to the underlying actin network [14].

In mice, global EHD2 deletion does not affect viability, in contrast with the embryonic lethality of a Notch1 or Dll4 knockout models [54, 55]. However, loss of EHD2 has been shown to increase the number of caveolae that were detached from the membrane and significantly reduced production of endothelial nitric oxide, potentially indicating a preference in endothelial function [56]; loss of Notch function has also been associated with reduced endothelial nitric oxide synthase activity [57]. In line with these observations, we observed that EHD2 knockdown resulted in an elevated number of detached caveolae in ECs. Of note, EHBP1 also increased the number of detached caveolae, suggesting a similar role for stabilizing caveolin pits at the plasma membrane. Our results also showed that EHBP1 does not colocalize with caveolae to the same extent as EHD2, where EHD2 demonstrated a near perfect association with caveolae. This observation may be indicative that EHBP1 is not recruited to all caveolae and may participate in binding to only a subpopulation of caveolin-mediated endocytic events, relating to its overall lower abundance.

Given EHBP1 had a greater impact on Notch signaling we preformed RNAseq to characterize the transcriptional landscape in its absence. A standout was differential expression of gene transcripts related to proteins that contained EGF repeats, such as Notch and Dll4. In one view this is not surprising, given proteins that contain EGF repeats are typically plasma membrane bound receptors or ligands that require endocytosis for general function, such as removal from the plasma membrane, recycling, etc. However, we were surprised at how narrow the list of differentially regulated genes was, with approximately 45% of hits relating to EGF repeat-containing proteins. We interpret these data as evidence that EHBP1 may play a very selective role in endocytosis, assisting internalization of proteins that require a robust stabilization by anchoring endocytic pits to the actin cytoskeleton, also support the notion that EHBP1 may be acting as specialized adaptor compared with EHD2 that is more ubiquitous to caveolae.

In aggregate, our results characterize EHBP1 and EHD2’s role in Dll4 caveolin-mediated endocytosis. Our analysis uncovered two major findings: 1) Dll4 uses a non-clathrin-mediated endocytic program; and 2) EHBP1 and EHD2 are required for Dll4 internalization during Notch receptor engagement. These results raise questions pertaining to the role of other endocytic proteins in the basal- and/or bound-state of Dll4. With regard to EHBP1 and EHD2, what is the precise pulling force contribution required for Dll4 endocytosis and how much is controlled by anchoring to the cytoskeleton is not known. It is also interesting to speculate how the caveolar machinery may interface with modifiers of Dll4, such as fringe proteins known to glycosylate Dll4’s extracellular domain. Overall, we believe that in addition to transcriptional regulation of Dll4/Notch proteins, the endocytic machinery involved in their signaling may add yet another level of regulation important for blood vessel development.

## ACKNOWLEDGEMENTS

Work was supported by funding from the National Heart Lung Blood Institute (Grant 1R56HL148450-01, R00HL124311) (A.M.W, C.R.F, and E.J.K). We like to thank Jennifer Bourne and the Electron Microscopy Center at the University of Colorado Anschutz Medical Campus for assistance with transmission electron micrograph collection. We would like to thank Cedric Asensio for critical reading of the manuscript.

## CONTRIBUTIONS

A.M.W, H.K, K.M.L and E.J.K created zebrafish and cell line constructs. A.M.W and E.J.K conceived experiments. R.J. performed mouse retinal experiments. A.M.W and C.R.F performed fibrin bead experiments. J.M.W and A.M.W performed RNA-seq analysis. S.M.M performed EHD2 TEM analysis. A.M.W. and E.J.K wrote the manuscript.

## MATERIALS AND METHODS

All zebrafish used in this study were AB strain. Zebrafish housing and protocols were all approved by the Institutional Animal Care and Use Committee (IACUC, number 946125-1). Zebrafish embryos were raised in a 28°C incubator in 1xE3 for 5 days. From 5-10 days post fertilization, embryos were raised at 28°C incubator with live marine L-type rotifers. From 10 day to 30 days post fertilization, larvae were fed GEMMA Micro 150 (Skretting USA). After 30 days post fertilization, zebrafish were fed GEMMA Micro 300 (Skretting USA). Clutches were composed of approximately 50% female and 50% male, which possess no defining sex characteristics at this time point.

Tg(kdrl:GFP) strain previously published by Choi et al. [58]. Tg(fli:LifeAct-GFP) strain previously published by Hen et al. 2015 [32]. Tg(kdrl:mCherry) strain previously published by Proulx et al. 2010 [59]. Tg(kdrl:cre) strain previously published by Hübner, K., et al. [25]. Tg(cdh5:gal4FF) strain previously published by Bussmann et al. 2011 [35].

Tol2 transposase RNA was synthesized from pT3TS-Tol2 (Addgene, #31831) using the MEGAscript™ T3 Transcription Kit (Thermo Fisher Scientific, AM1338) and stored at −80°C at a dilution of 100 ng/µl. Injection mixture was prepared on ice containing 300 ng Tol2 transposase RNA and 500 ng recombinant plasmid and was brought to 10 µl total volume with 0.1% phenol-red (VWR, 470301-974) in water. 1-4 cell embryos were injected directly into cell with 2 pL injection mixture.

4-guide CRISPR/Cas9 targeted gene KO was performed as outlined by Wu et al 2018 [21]. In brief, 4 single guide RNA templates fused to a scaffold were synthesized for each target gene using HiScribe™ SP6 RNA Synthesis Kit (New England BioLabs, E2070S). Injection mixture was prepared on ice containing 5 µM Cas9 (PNA Bio, CP02), 1 µg/µL sgRNA, and brought to 6 µL with 0.1% phenol-red in water. Cas9 and sgRNA guides were pre-complexed at 37°C for 5 mins. 1-4 cell embryos were injected directly into yolk with 2 pL injection mixture. Validation of 4 guide KO was validated in 3 fish by amplification of a single targeted guide site into a pME backbone (see table below) followed by Sanger sequencing by QuintaraBio.

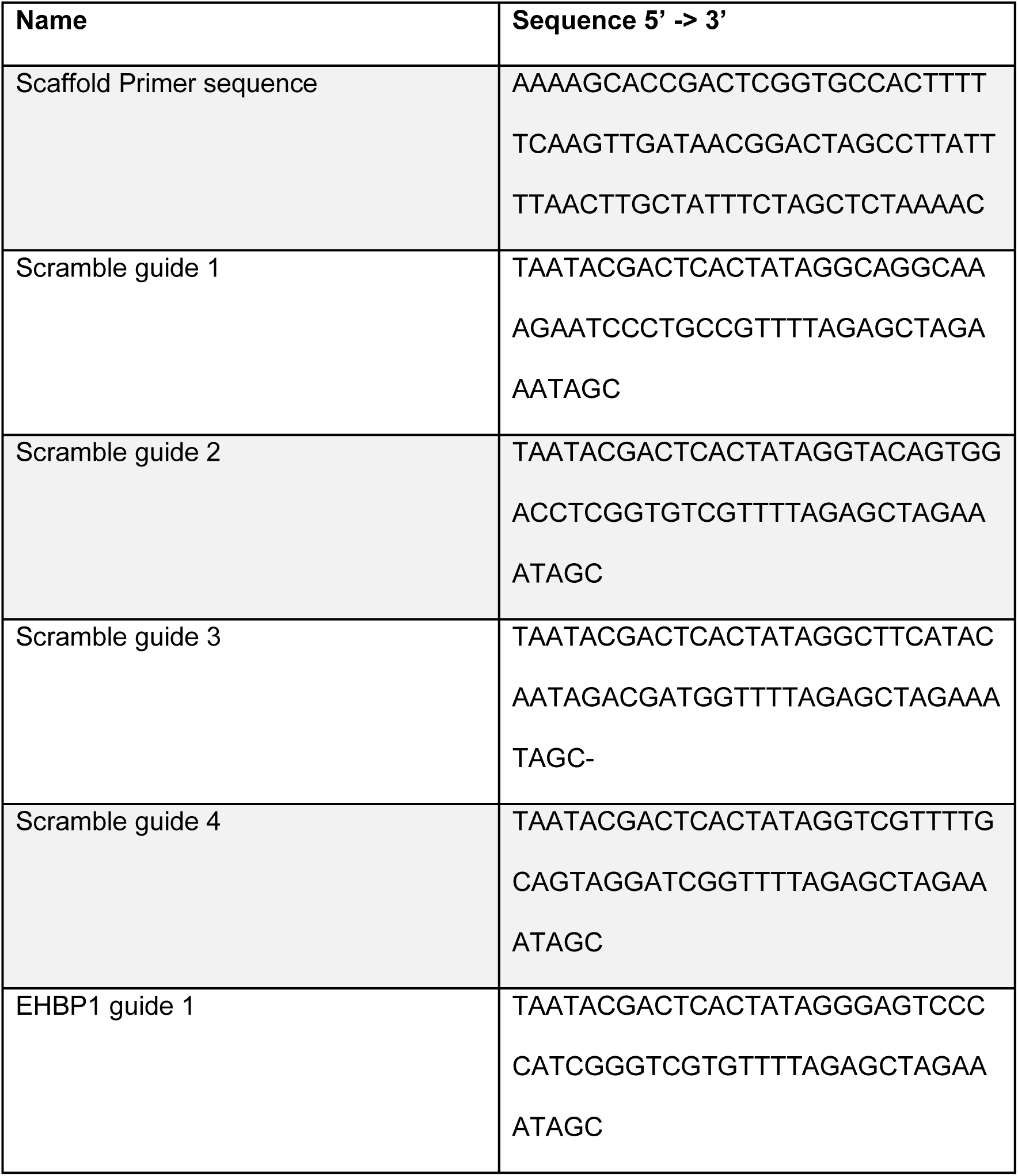
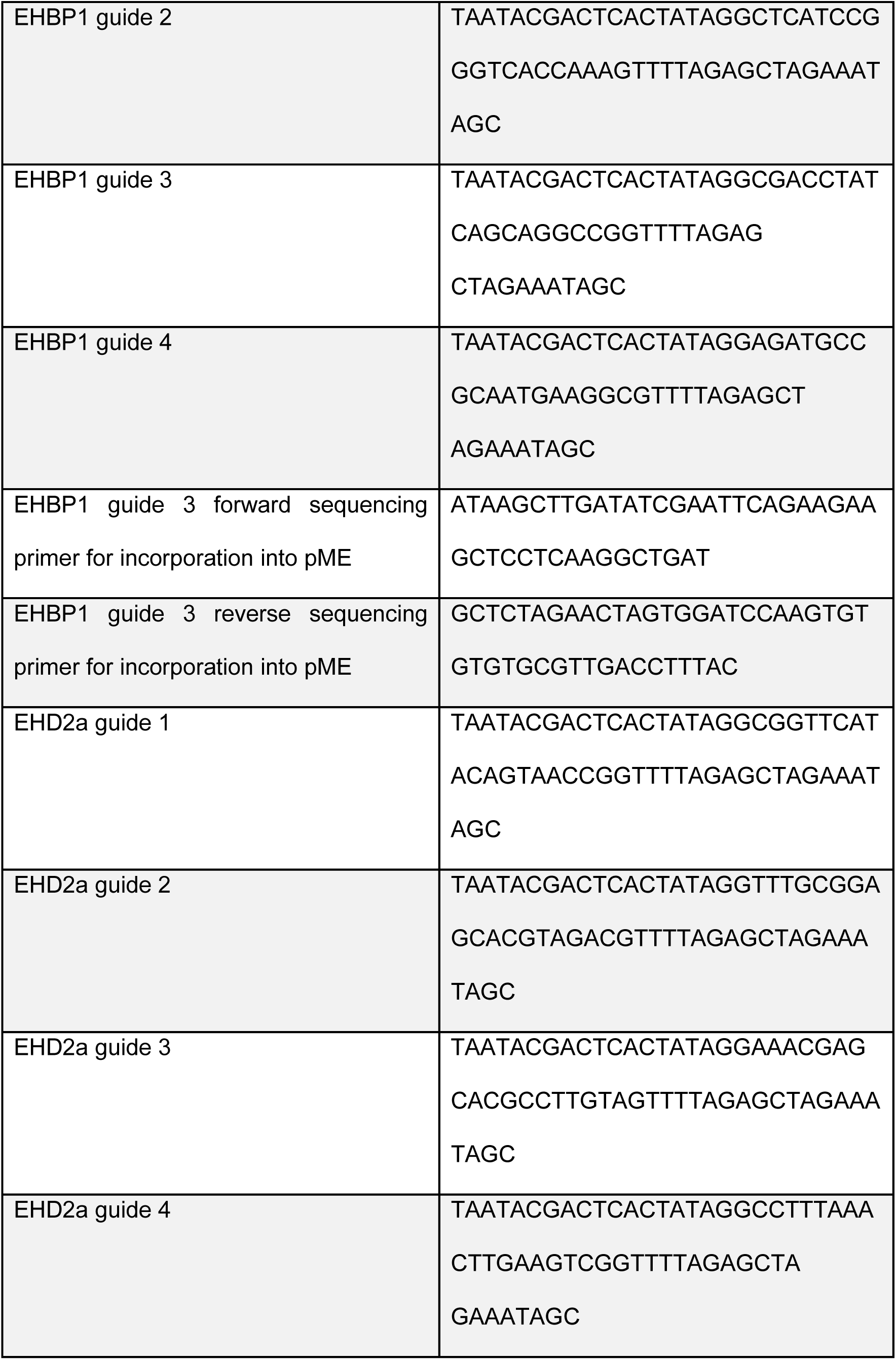
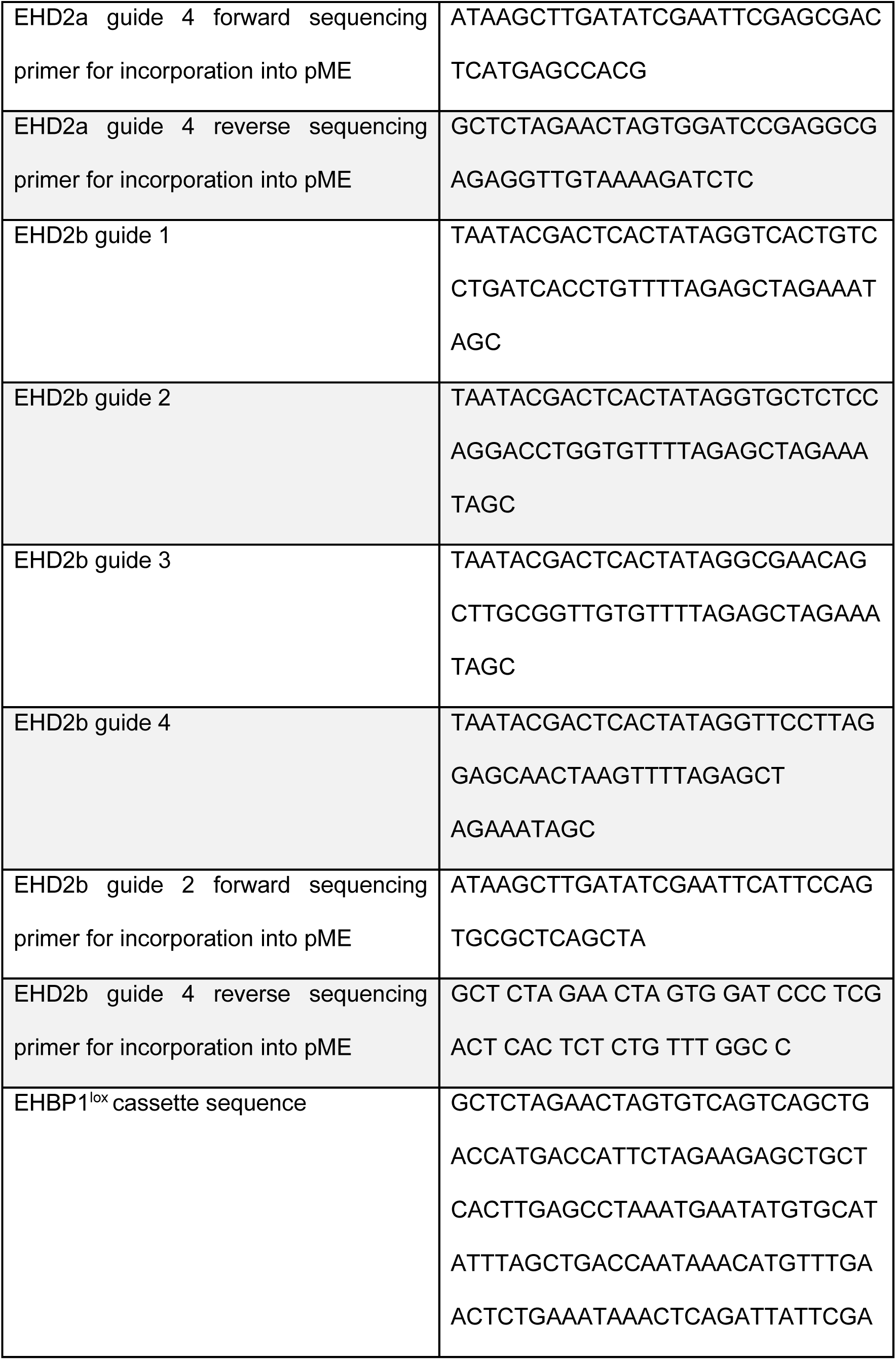
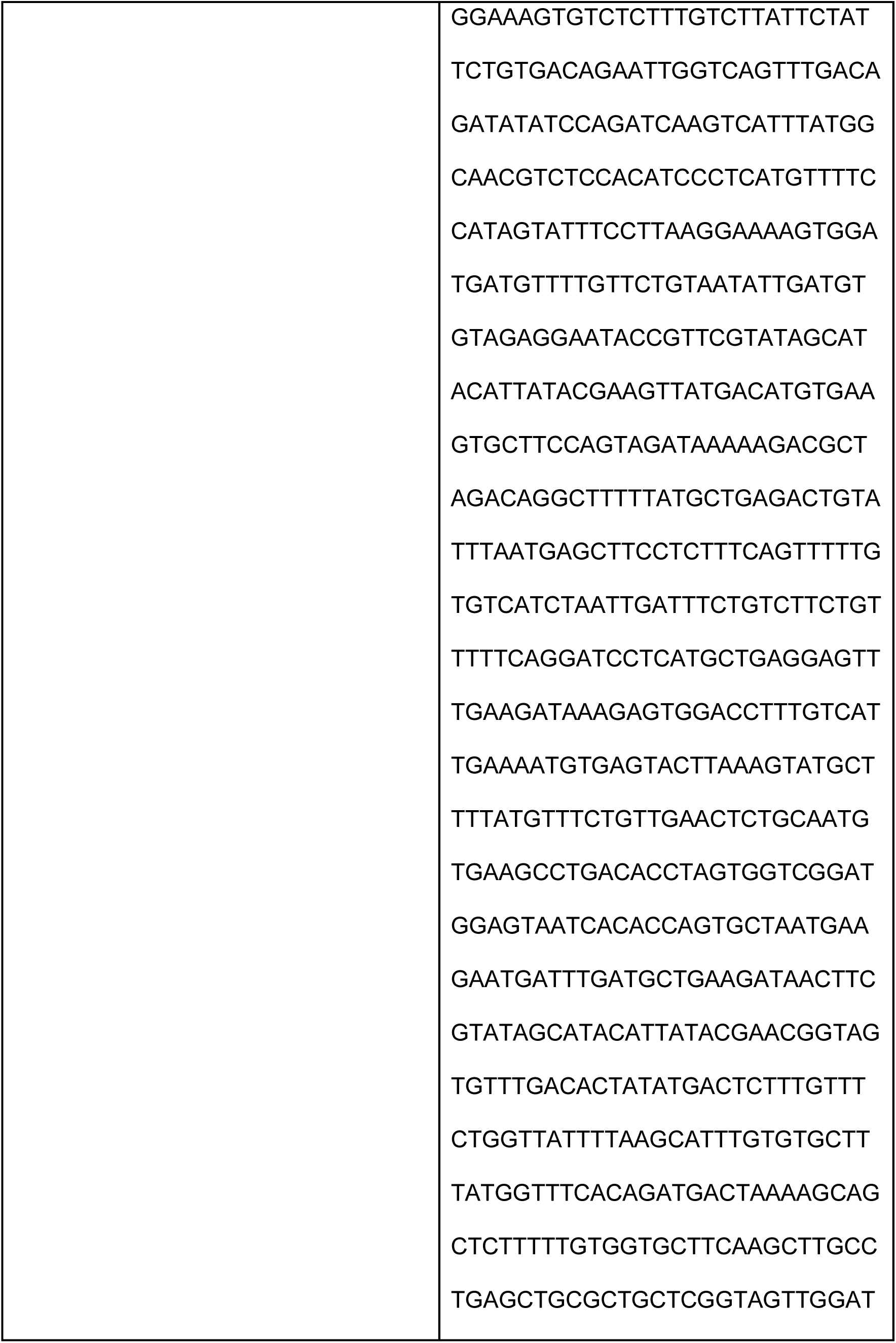
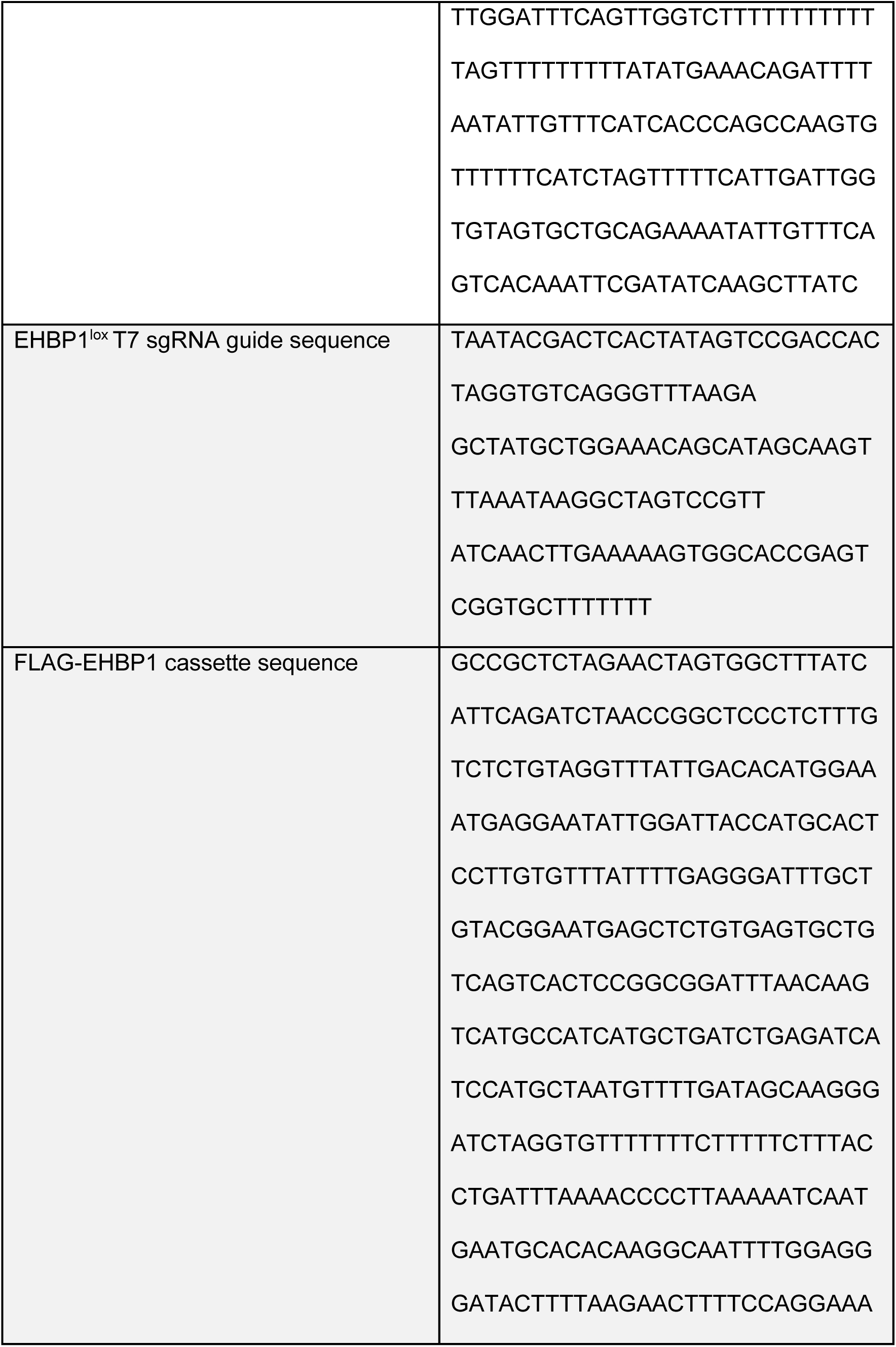
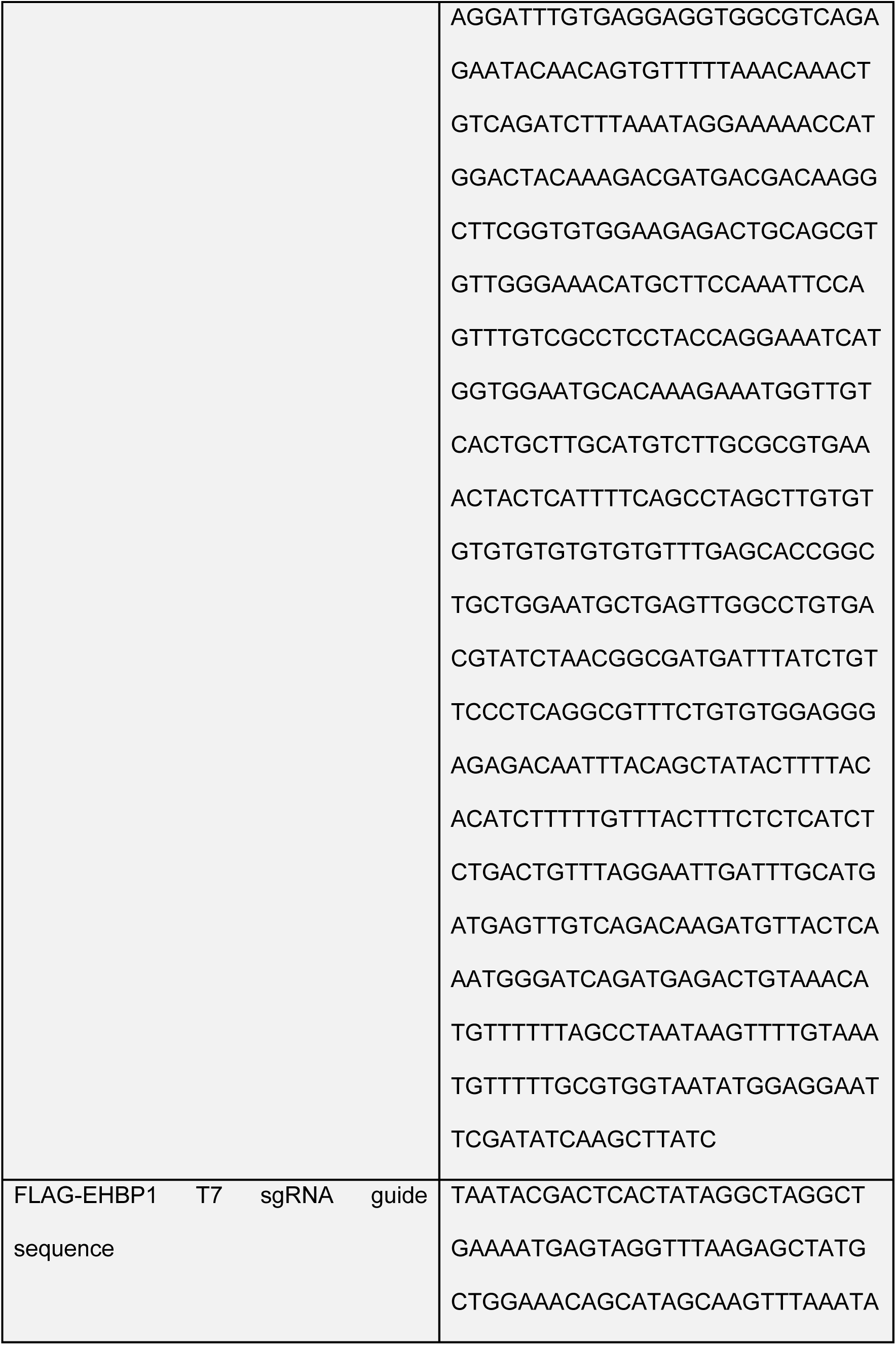
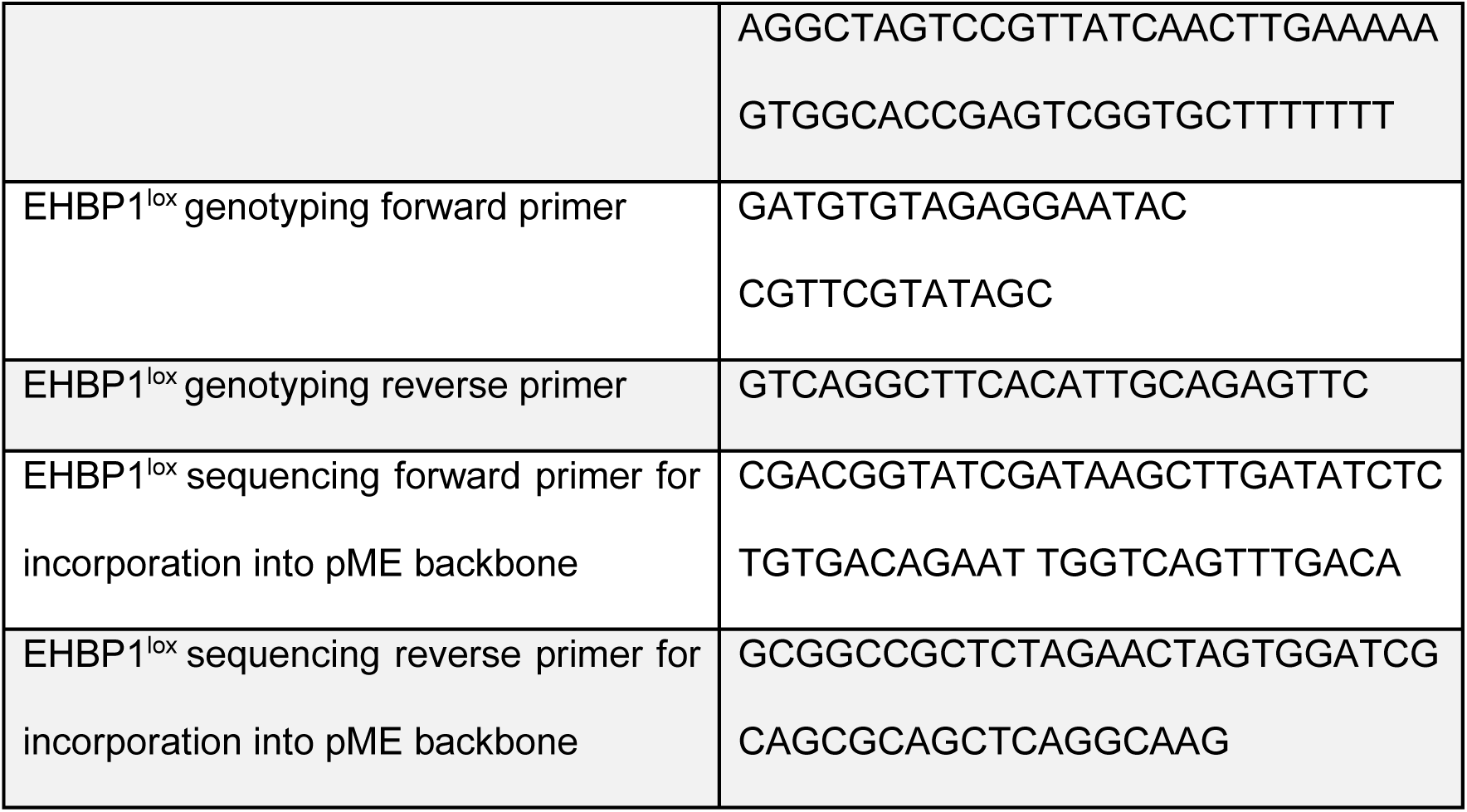

### In Situ Hybridization

In situ hybridization performed as outlined by Thisse and Thisse. 2007 [24]. DNA was primed from a zebrafish cDNA library using the primers below and inserted into a pME backbone containing both T7 and SP6 promoters via Gibson reaction. Antisense probes were converted to RNA from this template using the HiScribe™ SP6 RNA synthesis Kit (New England BioLabs, E2040S). Sense probes were converted to RNA from this template using the HiScribe™ T7 RNA synthesis Kit (New England BioLabs, E2070S). In each RNA synthesis reaction, UTP was substituted for DIG RNA labeling mix (UTP) (Sigma Aldrich, 11277073910). Probes were designed to be roughly 800 bp in size, which has shown to produce the most efficient labeling in zebrafish. Antisense probes were used to detect the transcript of interest, sense probes were used as a control to monitor overdevelopment of staining solution (225 µl Nitro Blue Tetrazolium [50 mg dissolved in 0.7 ml N,N-dimethylformamide anhydrous and 0.3 ml water], 50 ml Alkaline Tris Buffer [100 mM 1M Tris HCl pH 9.5, 50 mM MgCl_2_, 100 mM 5M NaCl, 0.1% Tween 20 20%], 175 µl 5-Bromo 4- Chloro 3-indolyl Phosphate [50 mg dissolved in 1 mL N,N-dimethylformamide anhydrous].

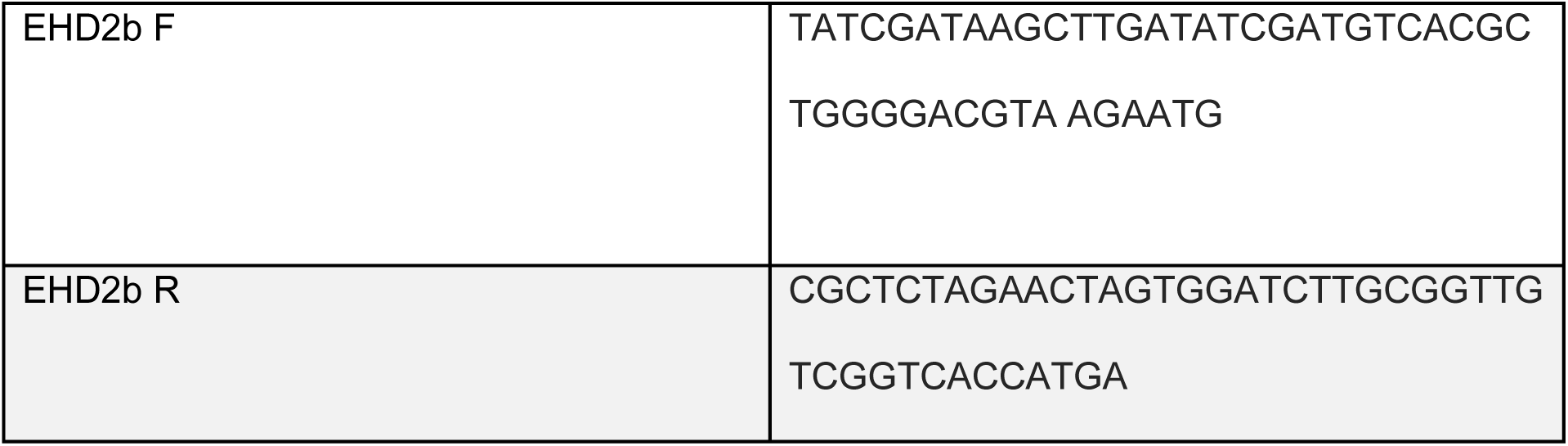

### Cell Culture

Glass-bottomed imaging dishes were exposed to UV light for 6 minutes and coated with Poly-D-Lysine (Thermo Fisher Scientific, A3890401) for a minimum of 20 minutes. Dishes were then coated with 15 µg/mL laminin mouse protein (Thermo Fisher Scientific, 23017015) overnight at 37°C. Cells were plated onto laminin coated dishes for 4-6 hours prior to imaging or fixation. 0.9 µM siRNA (Thermo Fisher Scientific, s225944, s26959, am4611) was introduced to primary human umbilical vein endothelial cells (HUVEC) using the Neon® transfection system (Thermo Fisher Scientific).

Adenovirus constructs (tagRFP-EHD2, tagRFP-EHBP1, Emerald-Clathrin) were created as previously described (He et al. 1998). In brief, constructs were introduced via Gibson Assembly® (New England BioLabs, E2611) into pShuttle-CMV (Addgene, #16403). PShuttle-CMV plasmids were then digested overnight with MssI (Thermo Fisher Scientific, IVGN0244) and purified using E.N.Z.A.® Gel Extraction Kit (Omega Bio-Tek, D2500-01). Linearized pShuttle-CMV plasmids were transformed into the final viral backbone using electrocompetent AdEasier-1 cells (Addgene, #16399). Successful incorporation of pShuttle-CMV construct into AdEasier-1 cells confirmed via digestion with PacI (Thermo Fisher Scientific, IVGN0184). 5000 ng plasmid was then digested at 37°C overnight, then 85°C for 10 minutes and transfected in a 3:1 polyethylenimine (PEI, Sigma Aldrich, 408747):DNA ratio into 70% confluent HEK 293A cells (Thermo Fisher Scientific, R70507) in a T-25 flask (Genesee Scientific, 25-207).

Over the course of approximately 2-4 weeks, fluorescent cells became swollen and burst or budded off of the plate. Once approximately 50% of the cells had lifted off of the plate, cells were scraped off and spun down at 2000 rpm for 5 minutes in a 15 mL conical tube. The supernatant was aspirated, and cells were resuspended in 1 mL DPBS (Genesee Scientific, 25-508B). Cells were then lysed by 3 consecutive quick freeze-thaw cycles in liquid nitrogen, spun down for 5 minutes at 2000 rpm, and supernatant was added to 2 70% confluent T-75 flasks (Genesee Scientific, 25-209). Propagation continued and collection repeated for infection of 10 15 cm dishes. After collection and 4 freeze thaw cycles of virus collected from 10 15 cm dishes, 8 mL viral supernatant was collected and combined with 4.4 g CsCl (Sigma Aldrich, 289329) in 10 mL DPBS. Solution was overlaid with mineral oil and spun at 32,000 rpm at 10°C for 18 hours. Viral fraction was collected with a syringe and stored in a 1:1 ratio with a storage buffer containing 10 mM Tris, pH 8.0, 100 mM NaCl, 0.1 percent BSA, and 50% glycerol. Primary Human Umbilical Vein Endothelial Cells (Promocell, C-12253) were treated with virus for 16 hours at a 1/1000 final dilution in all cell culture experiments.

Fibrin Bead Assay performed according to protocol as outlined by Nakatsu et al. 2007.

### pHrodo internalization assay

PHrodo™ iFL Red STP Ester (Thermo Fisher Scientific, P36010) was brought to 10 mM with DMSO. On the day of antibody labeling, pHrodo™ iFL Red STP Ester was diluted to 2 mM in DMSO (VWR Life Science, 97063-136). Antibody was brought up in DPBS (2 mg/mL final concentration for Dll4 polyclonal antibody [Thermo Fisher Scientific, PA5-46974]; 1 mg/mL final concentration for recombinant human Notch-1 Fc Chimera [R&D Systems, P46531]) and added to 1/10 volume 1 M sodium bicarbonate, pH 8.5. 3.3 µL of 2 mM pHrodo™ dye was added to the antibody and allowed to react in the dark for 1 hour with gentle flicking every 15 minutes. While this reaction is occurring, Zeba™ Spin Desalting Column (Thermo Fisher Scientific, 89882) was washed 3 times with 300 µL DPBS at 1500 x g for 1 minute. After 1 hour, the labeled antibody solution was loaded into the desalting column and allowed to absorb. DPBS was overlaid on top of labeled antibody to bring total column load volume to 70 µL and spun at 1500 x g for 2 minutes. Flow-through stored at 4 °C.

For bead tethered pHrodo, 6 µL of pHrodo-conjugated antibody brought to 200 µL TBST and added to 10 µL of either Dynabeads™ Protein G (Thermo Fisher Scientific, 10004D) or Protein A Agarose Resin (Gold Biotechnology, P-400-5). Beads and antibody were incubated and rotated at room temperature for 10 minutes. Conjugated beads were washed 3 times with 200 µL TBST then stored in a final volume of 10 µL TBST at 4°C.

3 µL pHrodo labeled antibody and 1 µL Hoechst 33342 trihydrochloride, trihydrate (Thermo Fisher Scientific, H3570) was added to 70-80 % confluent HUVECs plated on laminin coated dishes in 1 mL media and incubated at 37°C for 10 minutes. After 10 minutes, the cells were washed 3 times with 1 mL DPBS and then placed in 2 mL media. 10 z-stack images were taken for each condition, this marks time point 0 minutes. After images were taken of each group, cells were returned to 37°C for 10 minutes and imaged again.

### Microscopy

Images were taken on a Nikon Eclipse Ti inverted microscope equipped with a CSU-X1 Yokogawa spinning disk field scanning confocal system and a Hamamatusu EM-CCD digital camera. Cell culture images were captured using a Nikon Plan Apo 60x NA 1.40 oil objective using Olympus type F immersion oil NA 1.518 (ThorLabs, MOIL-30). Fish images were taken using either Nikon Apo LWD 20x NA 0.95 or Nikon Apo LWD 40x NA 1.15 water objectives. For transmission electron microscopy images, primary HUVEC of specified treatment were fixed in 2% glutaraldehyde, 2% PFA, 0.2 M Cacodylate buffer and sent to Jennifer Bourne at the University of Colorado Anschutz Medical Campus.

### Immunohistochemistry

2D cell culture was fixed in 4% paraformaldehyde in DPBS for 10 minutes. Cells were then washed 3 times for 5 minutes in TBST and permeabilized in 0.1% Triton-X 100 for 10 minutes. Cells were then washed 3 times for 5 minutes and blocked for 1 hour in 2% bovine serum albumin (BSA). Primary antibodies were applied at specified dilutions in Key Resources Table (Appendix A) overnight. Cells were washed 3 times for 10 minutes in TBST and then moved to secondary for 2 hours at specified dilutions in Key Resources Table. Cells were washed again 3 times for 15 minutes in TBST before imaging.

### Western Blot

Primary HUVEC culture was trypsinized and lysed using Ripa buffer (20 mM Tris-HCl [pH 7.5], 150 mM NaCl, 1 mM Na_2_EDTA, 1 mM EGTA, 1% NP-40, 1% sodium deoxycholate, 2.5 mM sodium pyrophosphate, 1 mM β-glycerophosphate, 1 mM Na_3_VO_4_, 1 µg/mL leupeptin) containing 1x ProBlock™ Protease Inhibitor Cocktail −50, Plus EDTA (GoldBio, GB-334-20). Total concentration of protein in lysate was quantified using the Pierce™ BCA Protein Assay Kit (Thermo Fisher Scientific, 23225) measured at 562 nm and compared to a standard curve. 20-50 µg protein was prepared in 0.52 M SDS, 1.2 mM bromothymol blue, 58.6% glycerol, 75 mM Tris pH 6.8, and 0.17 M DTT. Samples were boiled for 10 minutes, then 35 µL was loaded in a 7-12% SDS gel and run at 170 V. Protein was then transferred to Immun-Blot PVDF Membrane (BioRad, 1620177) at 4°C, 100 V for 1 hour 10 minutes. Blots were blocked in 2% milk for 1 hour, then put in primary antibody at specified concentrations overnight. After 3 10-minute washes with TBST, secondary antibodies at specified concentrations were applied for 4 hours. After 3 additional TBST washes, blots were developed with ProSignal® Pico ECL Spray (Genesee Scientific, 20-300S).

### Pharmacological Treatment

DAPT (Sigma Aldrich, D5942) was applied to cells for 3 days at a final concentration of 5 µM. Dynasore hydrate (Sigma Aldrich, D7693) was applied to cells for 30 minutes at a final concentration of 100 µM. LY-411575 (Sigma Aldrich, SML0506) was diluted in egg water to a final dilution of 2 µM from 30-48 hpf. Latrunculin A (Sigma Aldrich, 428021-100UG) was applied to cells for 1 hour at a final concentration of 5 µM. Cytochalasin B (Sigma Aldrich, C6762-5MG) was applied to cells for 1 hour at a final concentration of 10 µM. Methyl-β-Cyclodextrin (Sigma Aldrich, M7439-1G) was applied to cell for 10 minutes at a final concentration of 10 mM.

### RNA-Sequencing

RNA collected from siRNA treated HUVEC was collected in triplicate and sent to Novogene for RNA-sequencing. Sequence files received from Novogene were processed using the Bioconductor package in R-Studio as described by Love et al. [60] using Homo Sapiens GRCh38 version 98 as a reference genome.

### Quantification and Statistical Analysis

Ectopic vessels were defined by a sprout emerging out of or separate from the defined ISV and quantified in 48 hpf embryos expressing tg(kdrl:GFP). Number of filopodia per ISV was defined by filopodial extensions (observed with zebrafish line tg(fli:LifeAct-GFP)) emerging from a fully formed ISV that has connected with the dorsal longitudinal anastomotic vessel. RT-PCR was quantified using the gel analysis function in Fiji image analysis software (Schindelin et al., 2012). In sum, rectangular sections were drawn around individual lanes in gray-scale, high-quality gel image using the pathway *Analyze > Gel > Select First Lane, Analyze > Gel > Select Next Lane*. After all lanes are selected, the pathway *Analyze > Gel > Plot Lanes* was used. The peaks of each lane were then segmented using the Straight Line selection tool, and highlighted with the Wand tool. Selection of the area inside the peak generates a Results window with the area and percent of each peak. The percent value of each sample was divided by the percent value of the control to obtain a relative density. Relative densities of the gene of interest (e.g. Hey2) were divided by the relative density of the housekeeping gene (GAPDH) to obtain a final adjusted density value.

Cellular uptake of pHrodo-labeled antibody was also quantified using Fiji image analysis software. Stack files were z-stacked at maximum intensity, and each color channel was adjusted so that the background was zero. Each individual cell was outlined with the Freehand Selections tool. The color channels were then separated and any background fluorescence (488 channel) was subtracted from the pHrodo fluorescent intensity (561 channel) using pathway *Process > Image Calculator*. The Integrated Density of pHrodo fluorescent intensity within each cell boundary was then recorded for every cell at each time point. Each experiment was performed in duplicate.

Pearson’s coefficient was calculated using the Image J Plugin Just Another Colocalization Plugin (JACoP) [61]. All statistical analysis was performed in GraphPad Prism8. Comparisons between two conditions were made using a t-test, comparisons between multiple conditions were made using a One-Way ANOVA.

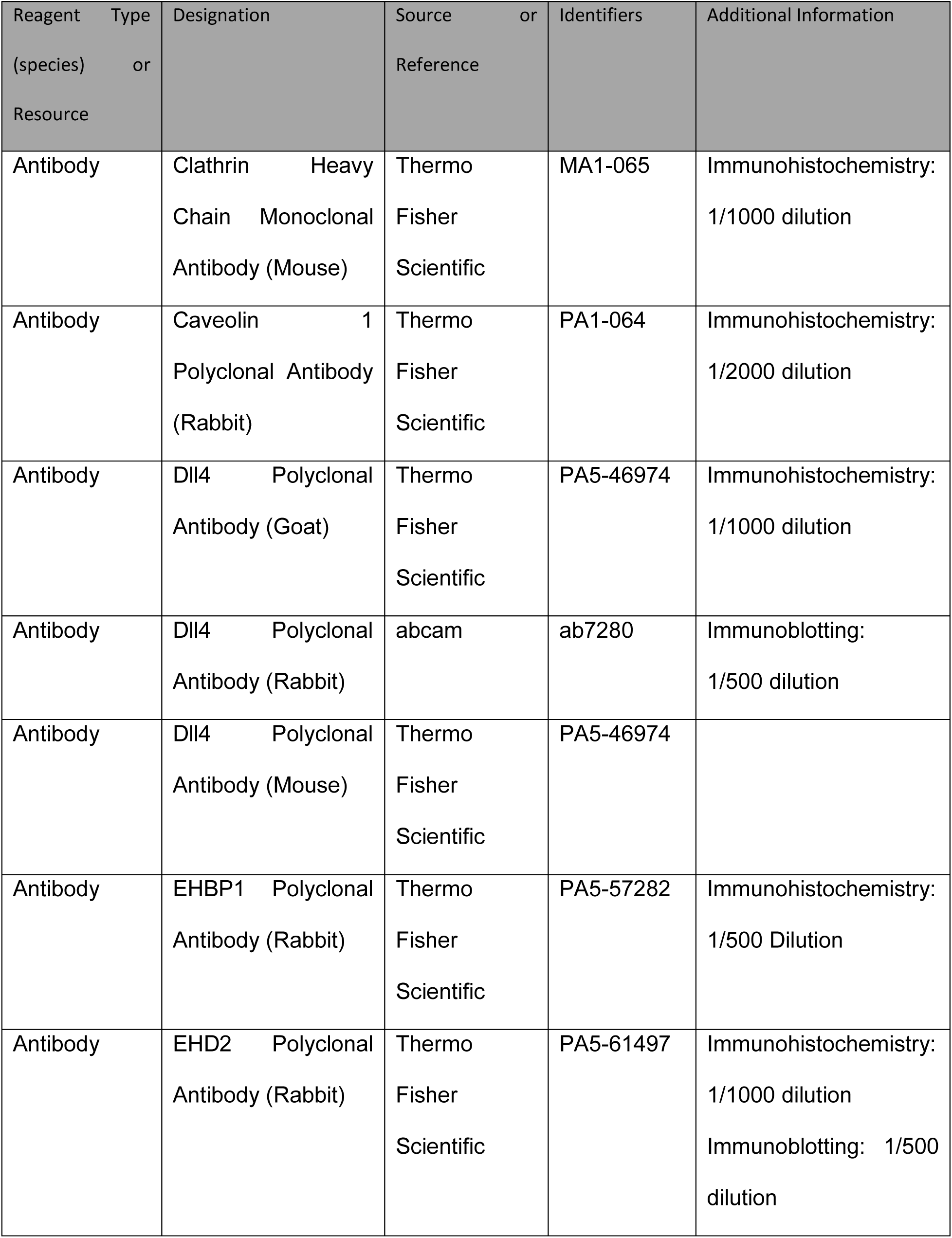
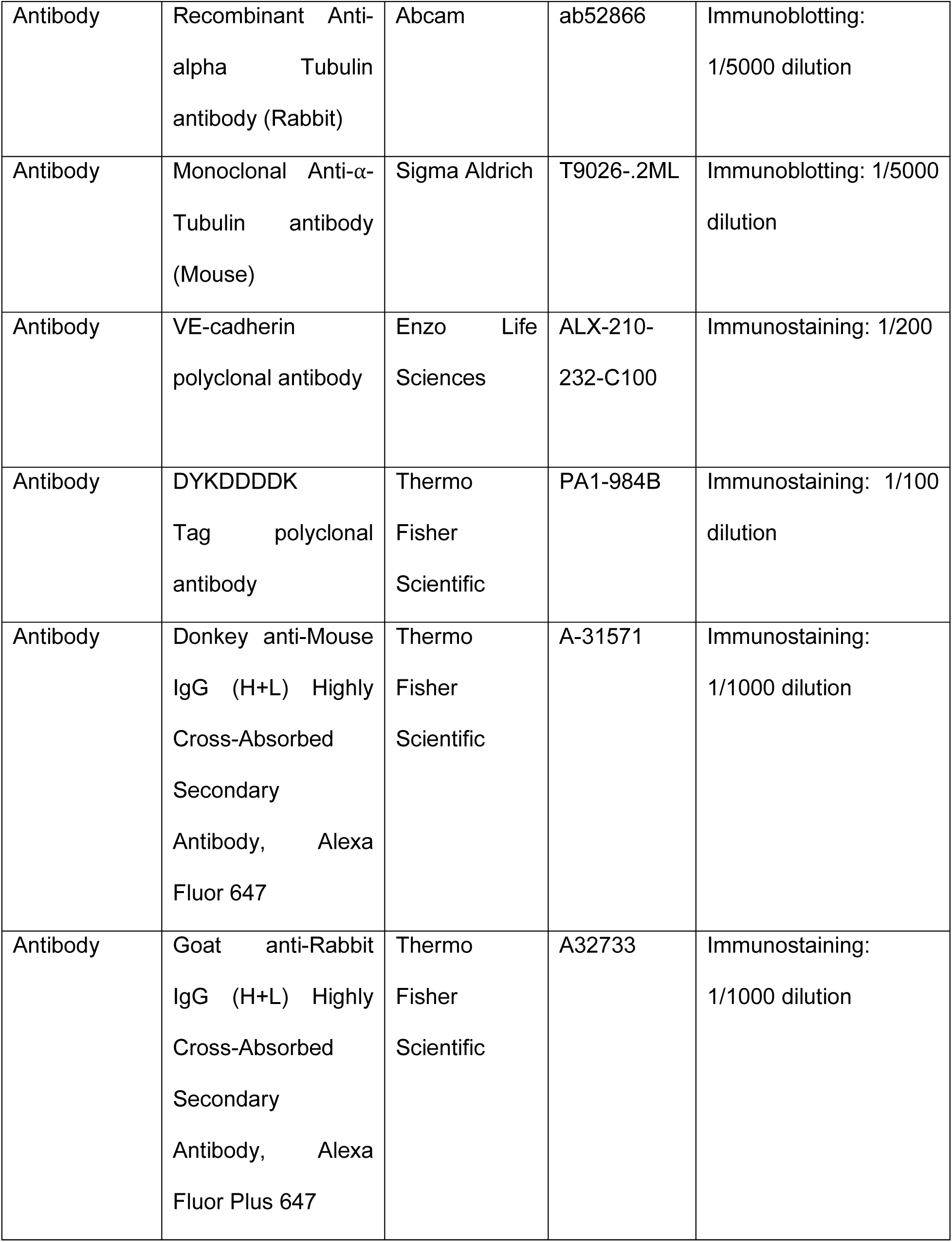
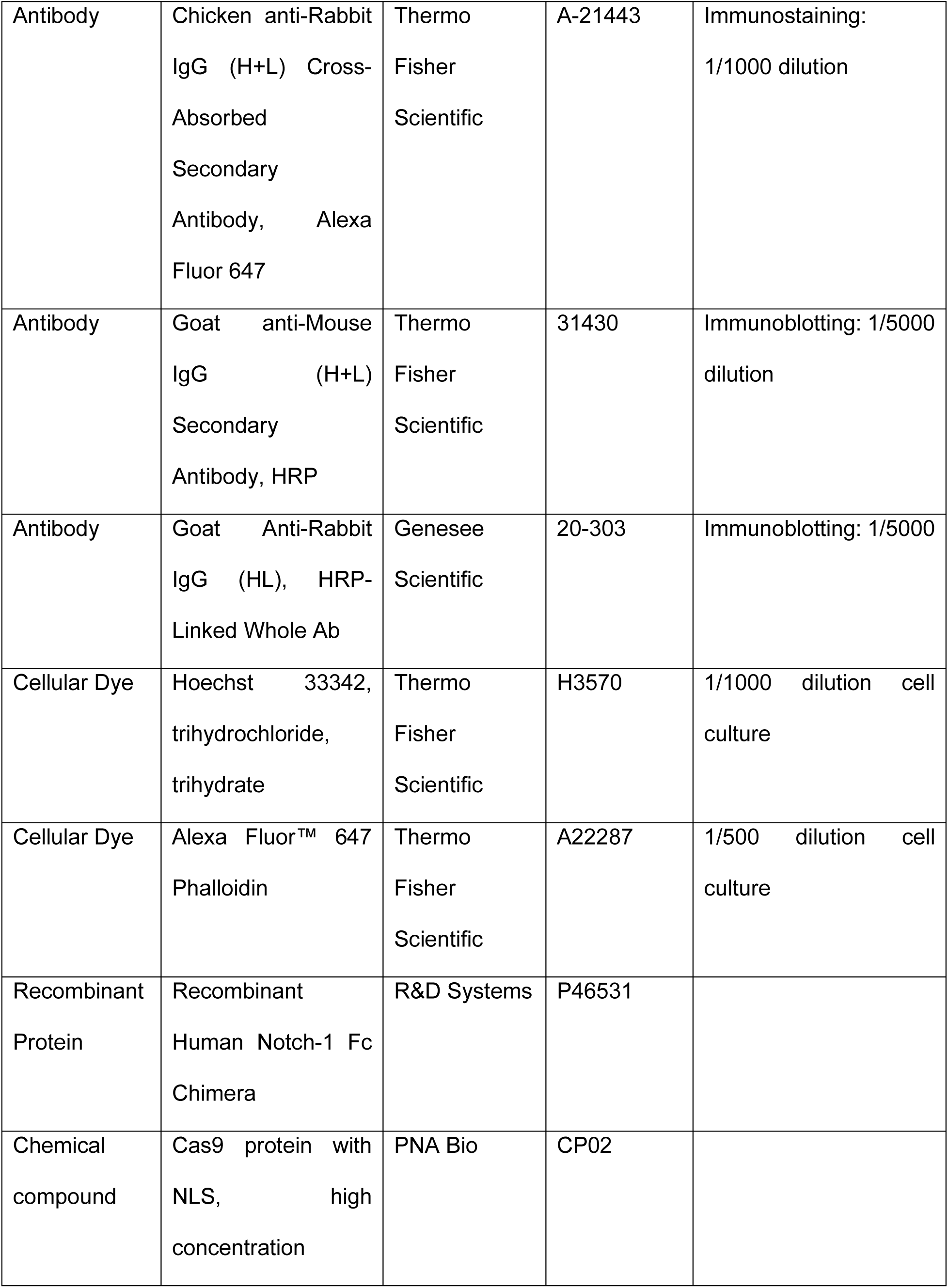
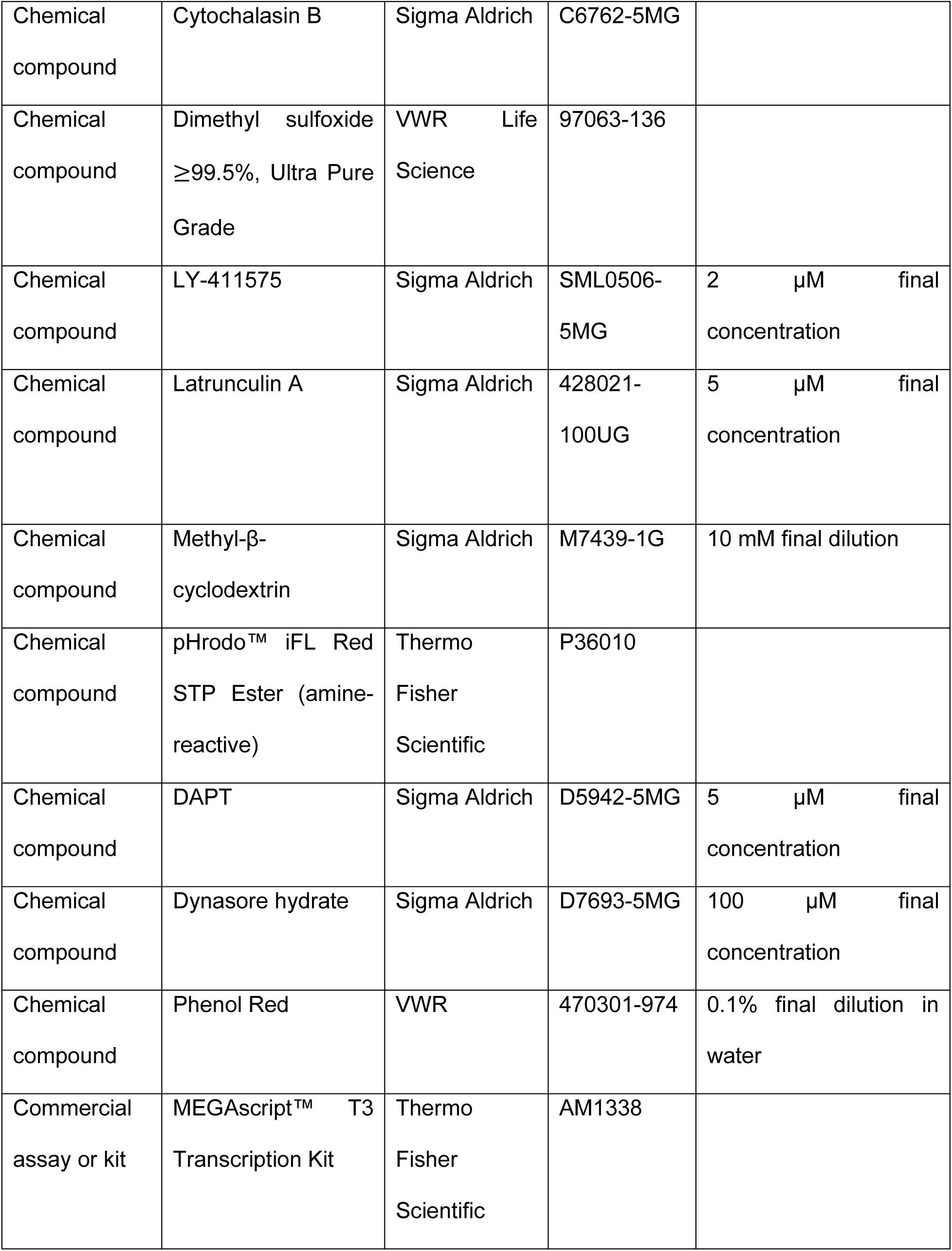
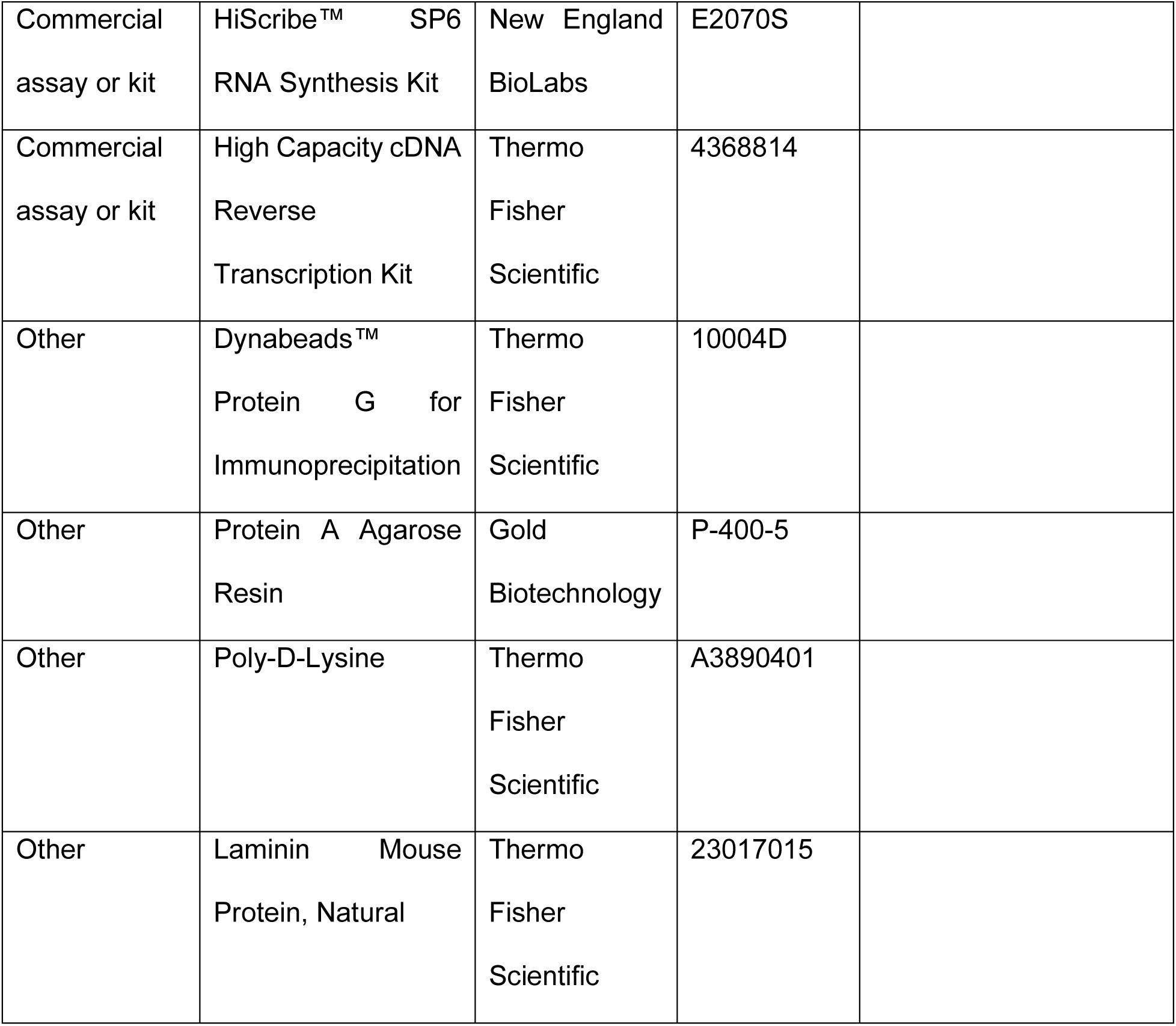

---

**Figure S1.**
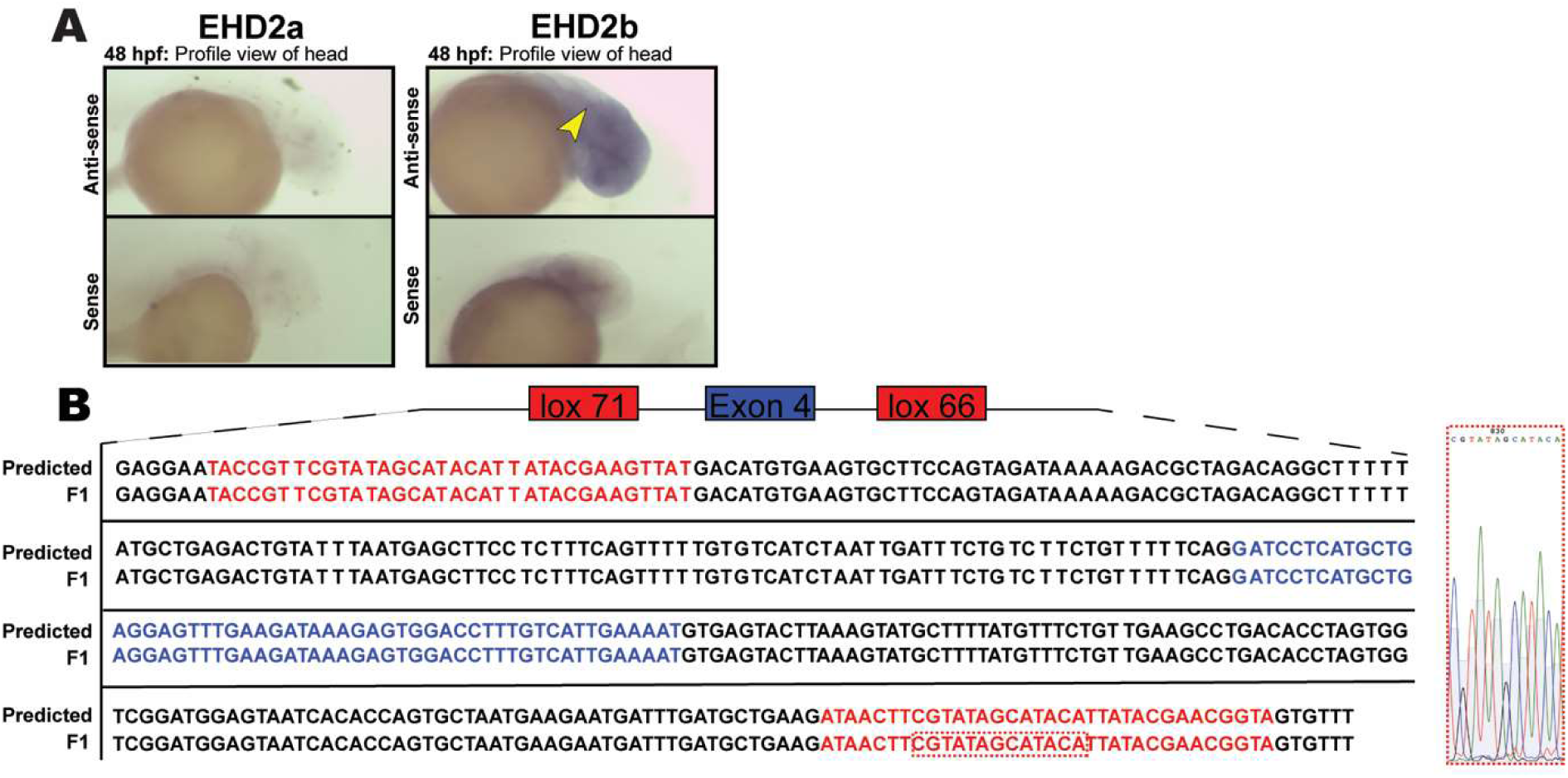
Confirmation of in vivo genetic tools. (A) In situ hybridization against either EHD2a or EHD2b. Yellow arrow points to expression along neural tube (C) Confirmation of in frame lox site incorporation into F1 generation EHBP1 floxed fish. Predicted sequence shown in first row, sequenced F1 fish shown in second row. Sanger sequencing peaks of small region in c-terminal lox site highlighted to the right.

**Figure S2.**
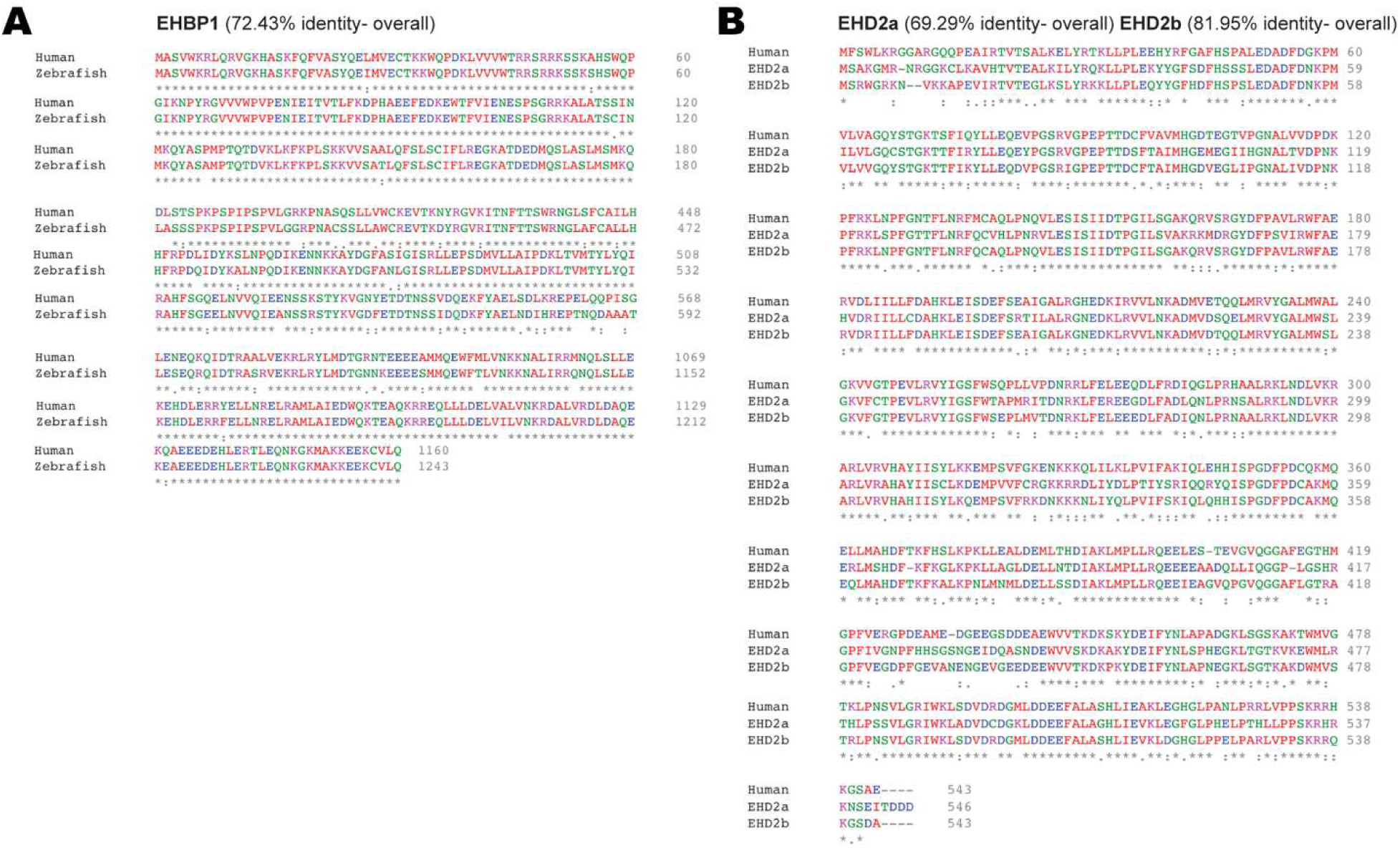
Comparison of EHBP1 and EHD2 amino acid identity between Human and Zebrafish. (A) Amino acid alignment of Human and Zebrafish EHBP1. (B) Amino acid alignment of Human EHD2 and zebrafish EHD2a or b.

**Figure S3.**
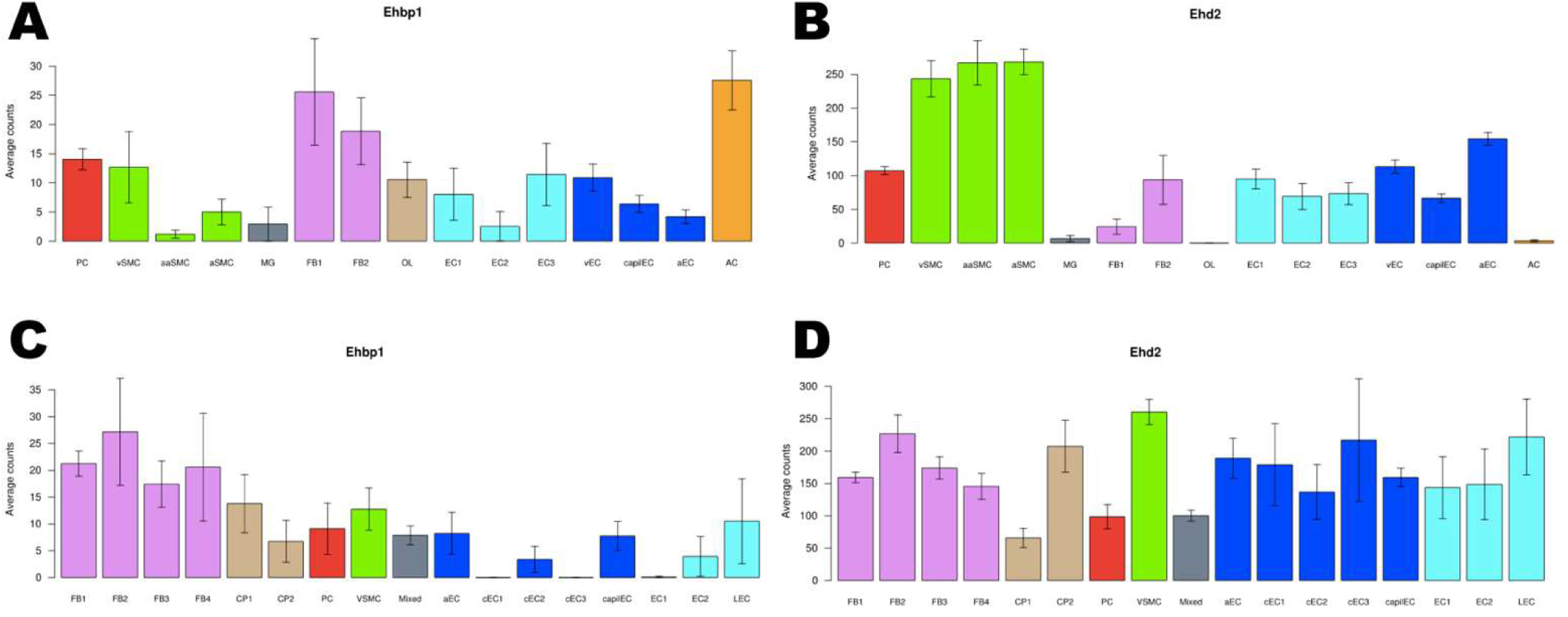
Single cell vascular expression data of EHBP1 and EHD2. (A,B) Brain data, average expression in each cluster. (C,D) Lung data, average expression in each cluster. [Brain data]: PC - Pericytes; SMC - Smooth muscle cells; MG - Microglia; FB - Vascular fibroblast-like cells; OL - Oligodendrocytes; EC - Endothelial cells; AC - Astrocytes; v - venous; capil - capillary; a - arterial; aa - arteriolar; 1,2,3- subtypes. [Lung data]: FB - Vascular fibroblast-like cells; CP - Cartilage perichondrium; PC - Pericytes; VSMC - Vascular smooth muscle cells; EC - Endothelial cells; capil - capillary; a - arterial; c - continuum; L - Lymphatic; 1,2,3,4 - subtypes.

**Figure S4.**
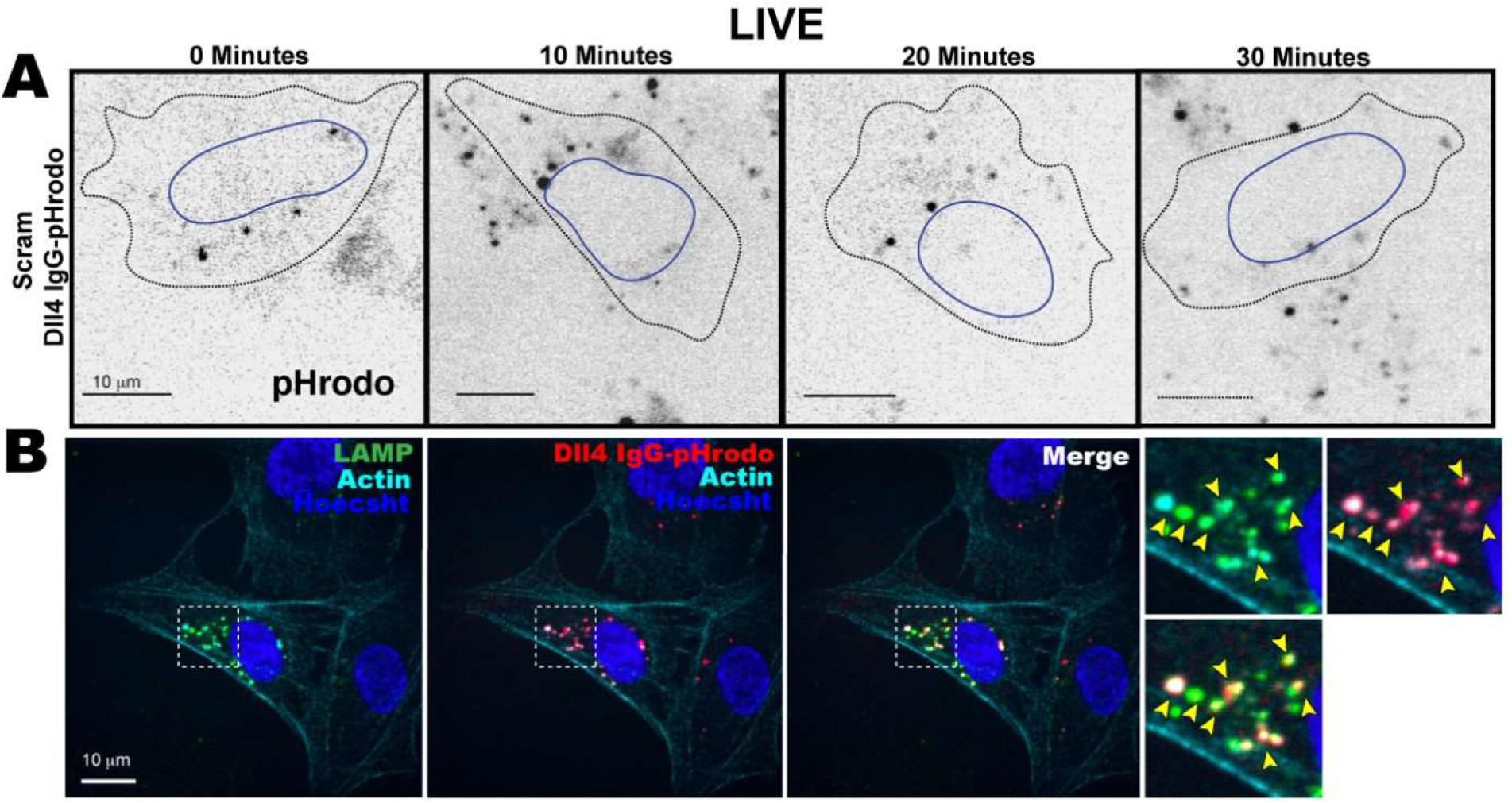
Dll4 IgG-pHrodo trafficks to lysosome. (A) Representative images of individual ECs internalizing pHrodo-Dll4 over time. pHrodo shown is black puncta. (B) Phrodo-treated ECs stained for lysosomal marker LAMP and actin. Yellow arrows show colocalization. ENSG00000038427 93.1004752 2.60E+00 0.51313274 5.06633197 4.06E-07 1.61E-04 VCAN

**Table S1.**
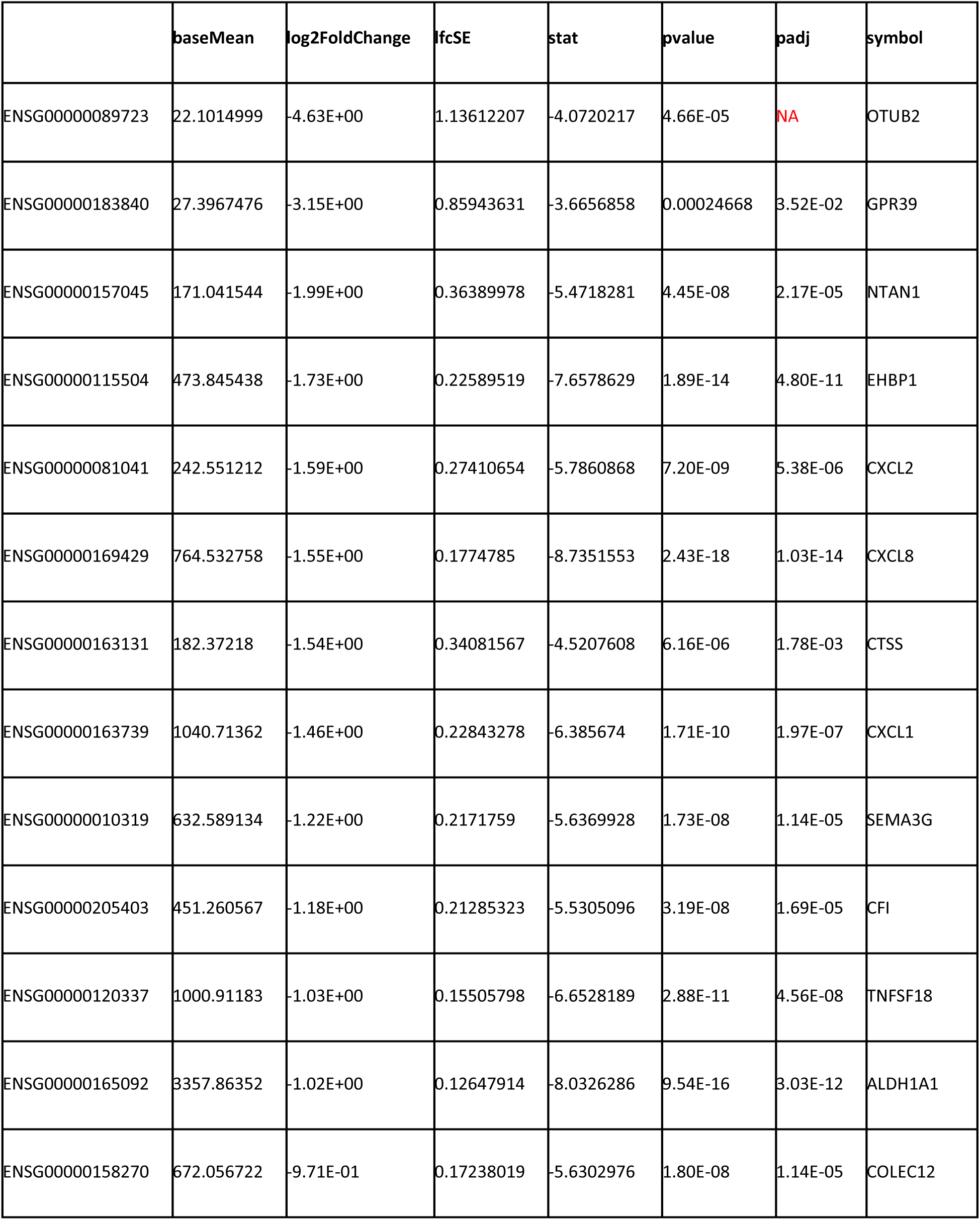
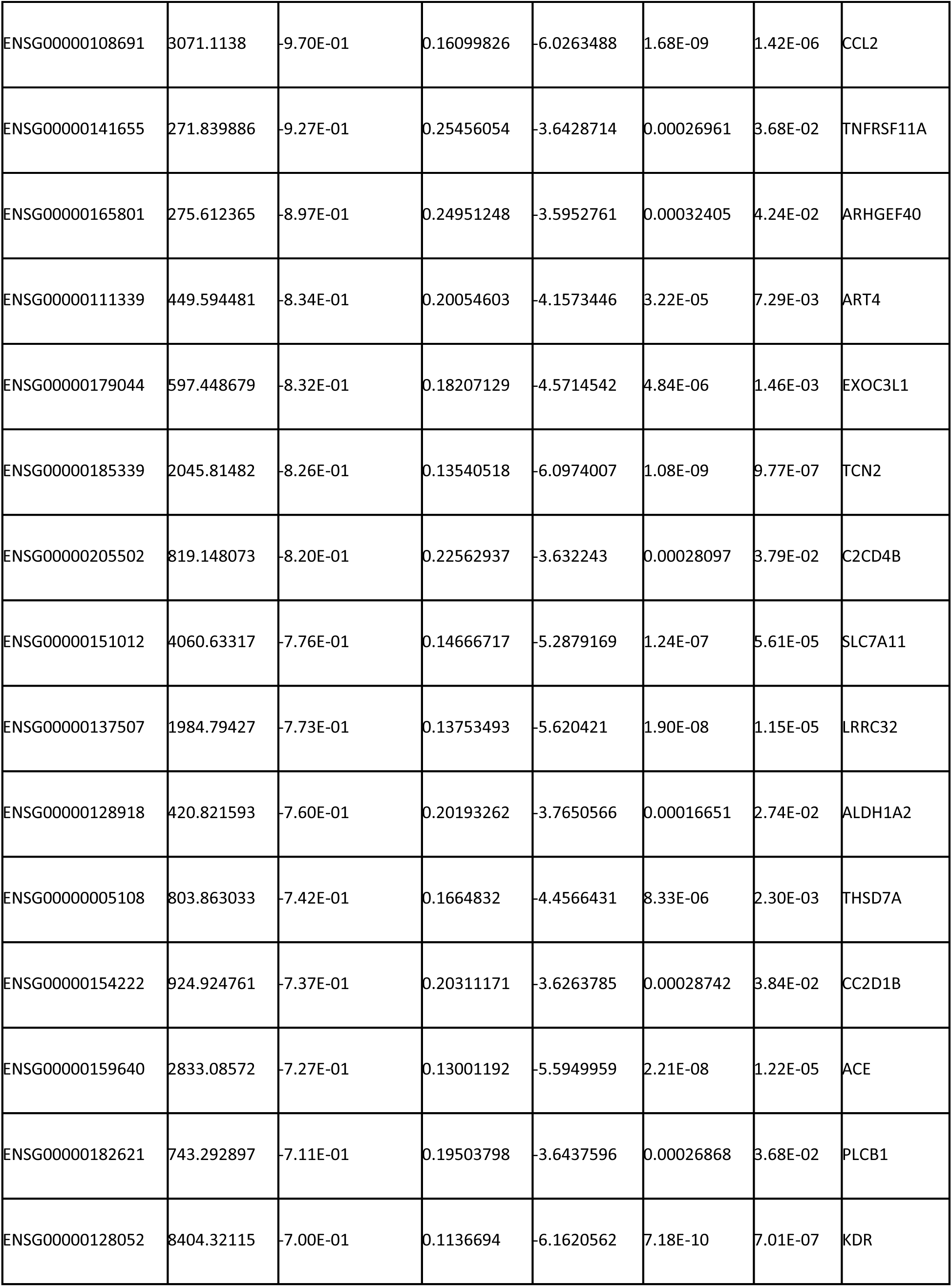
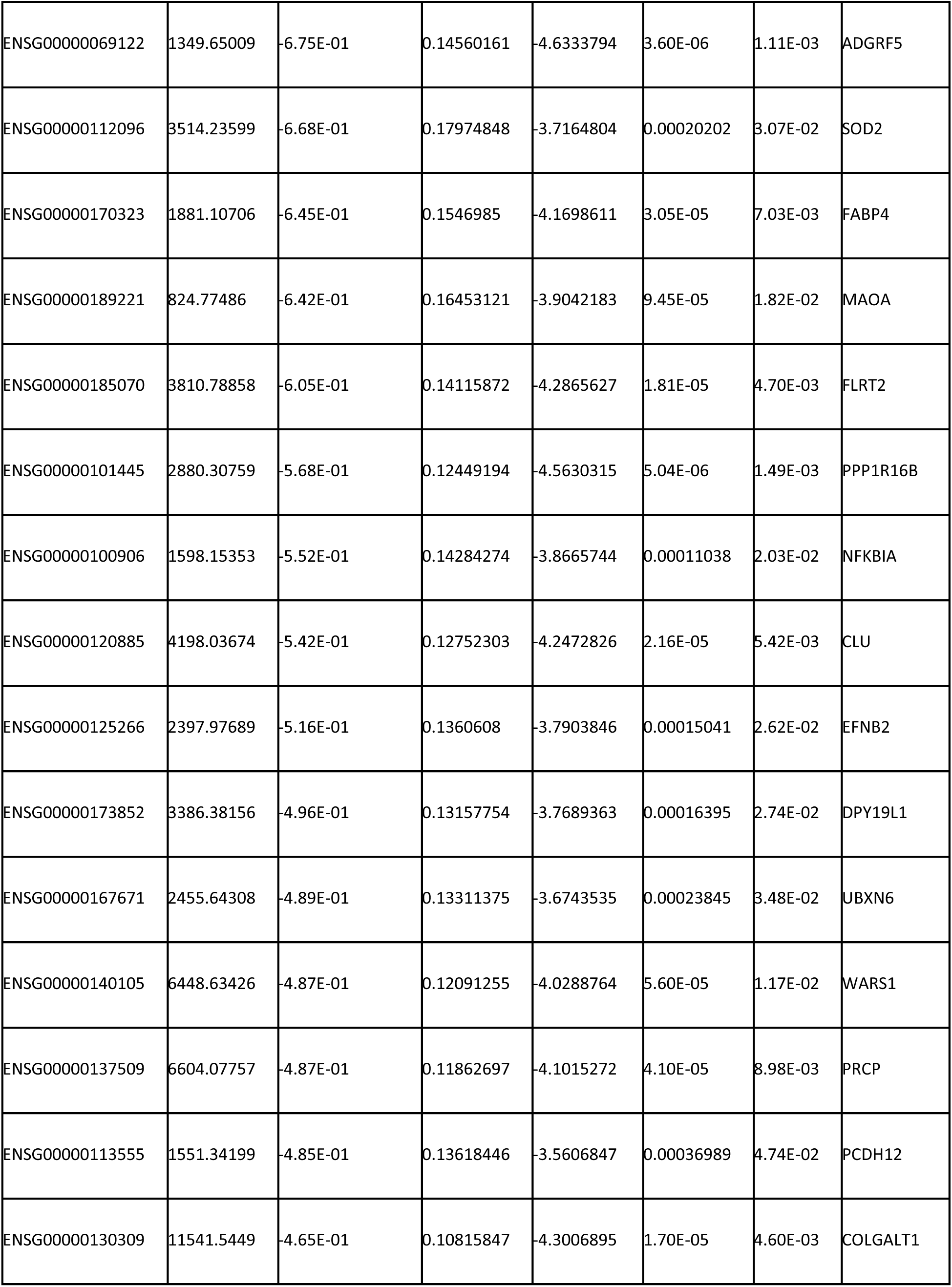
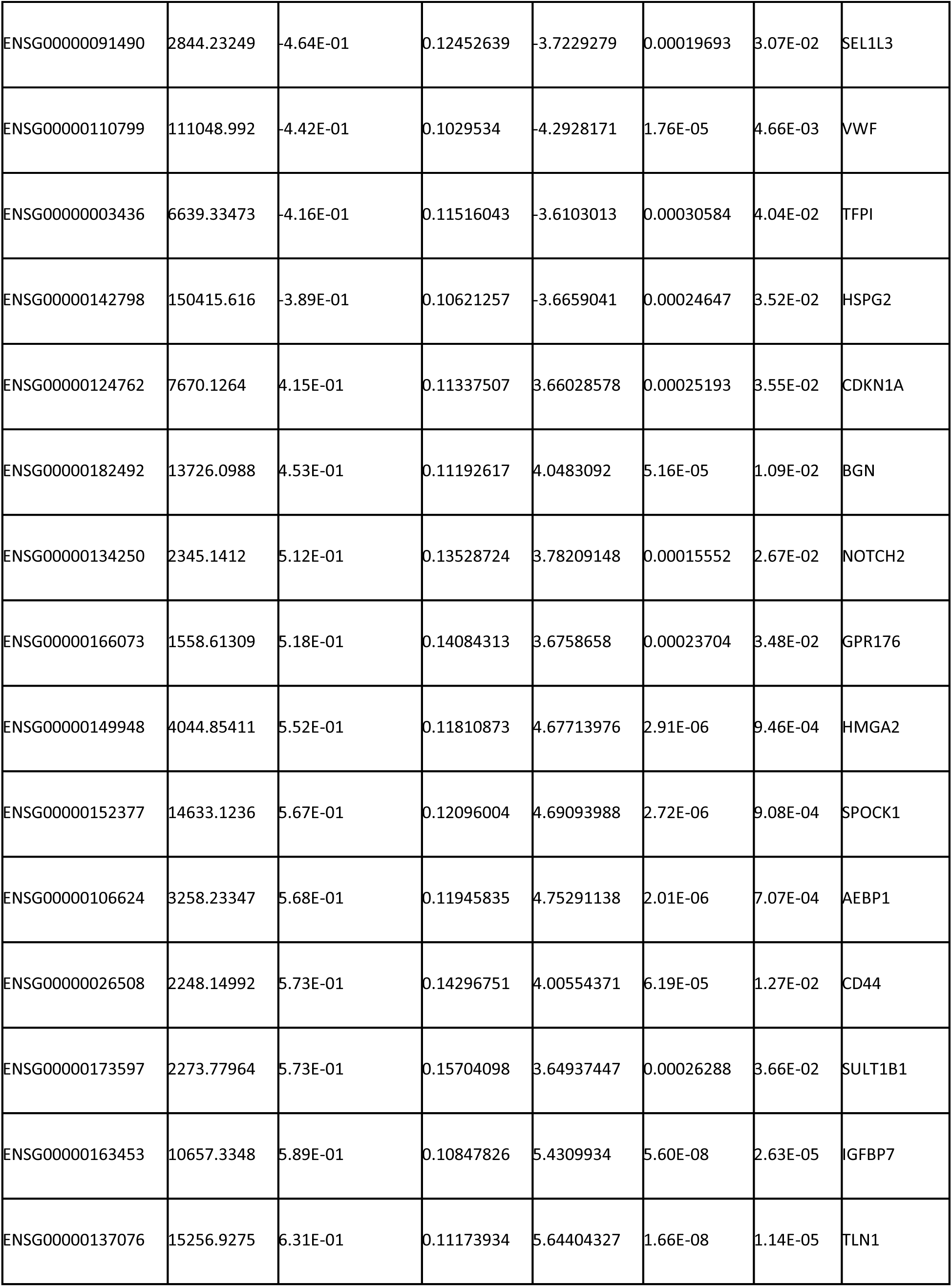
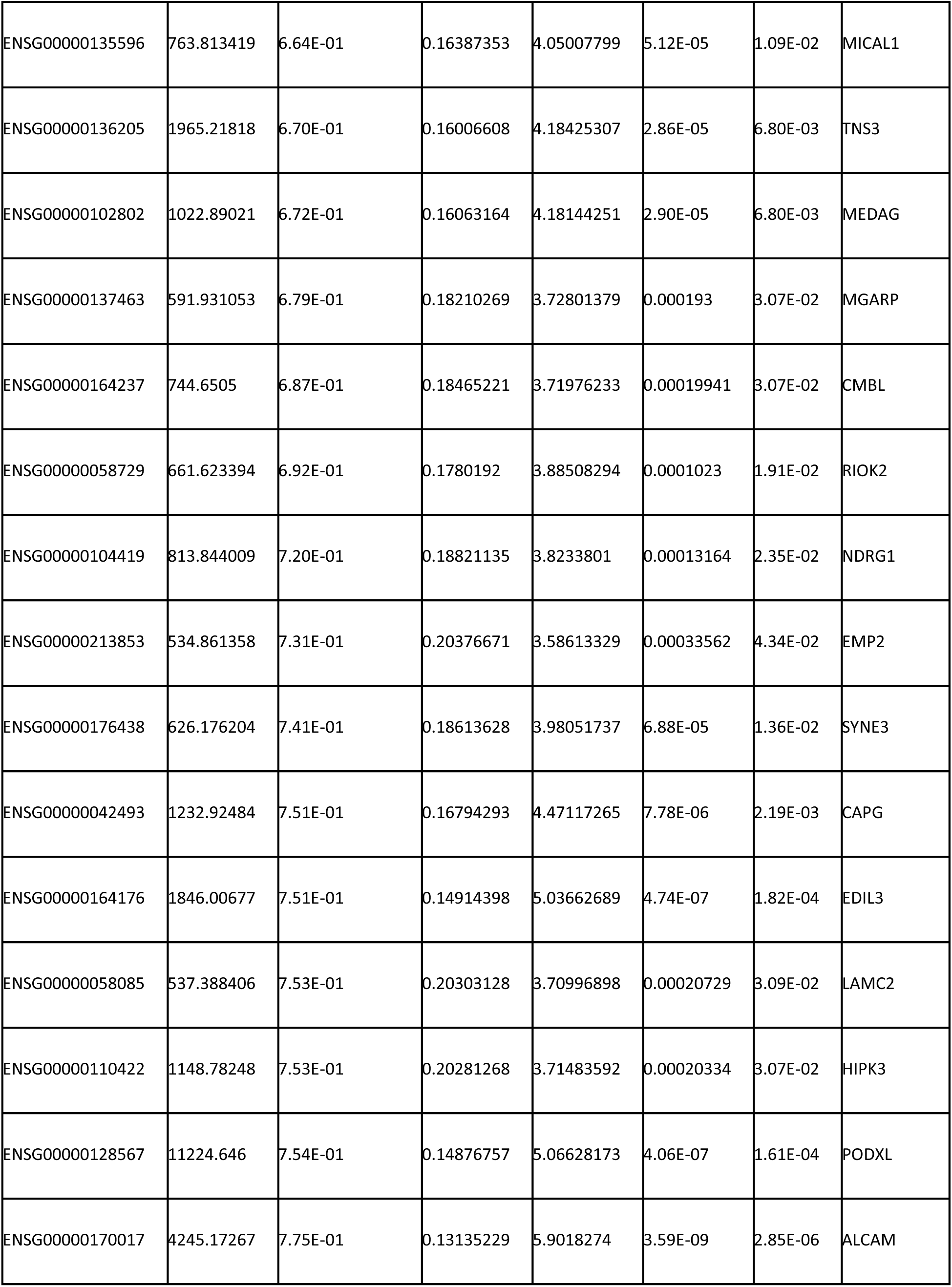
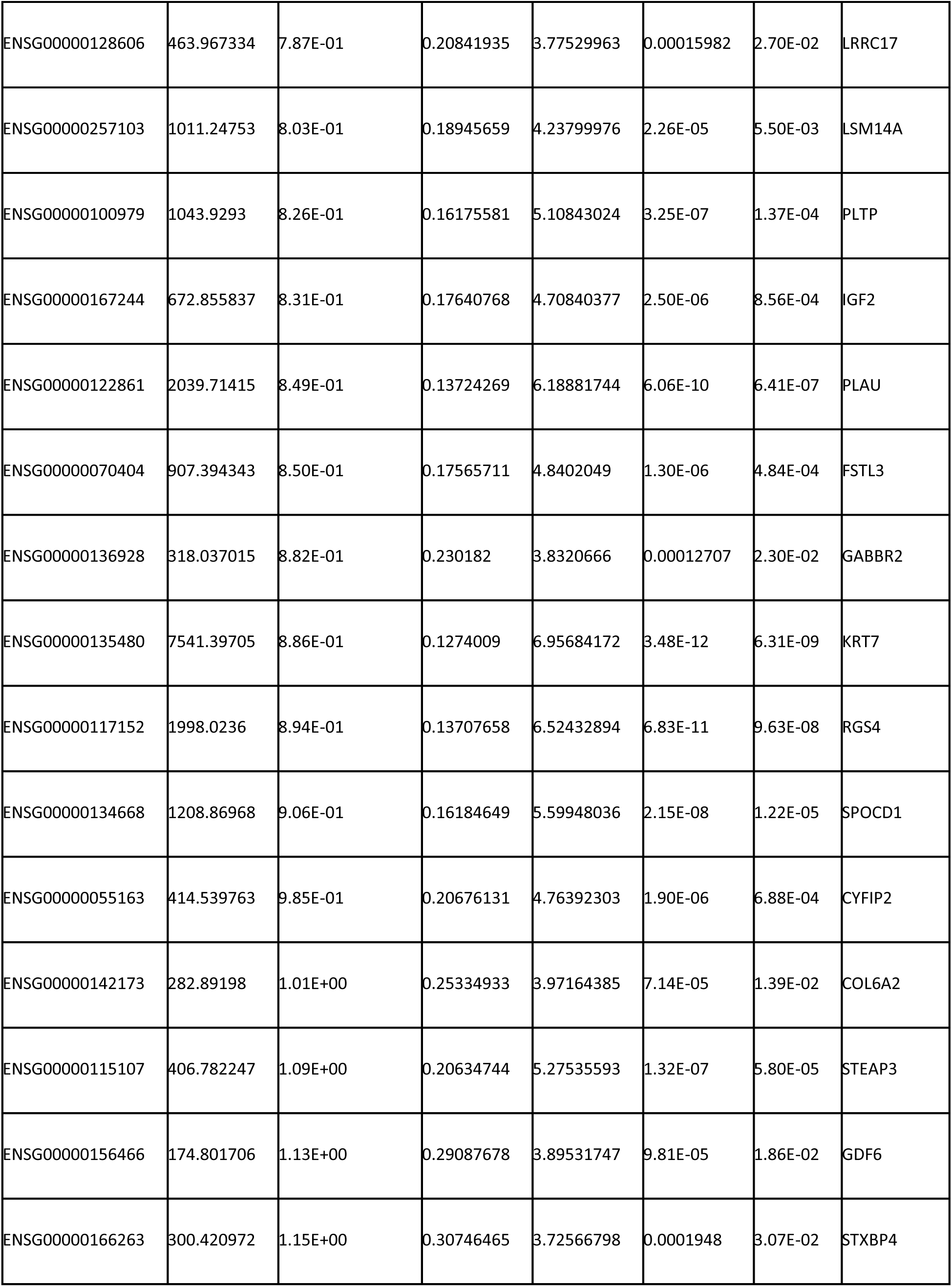
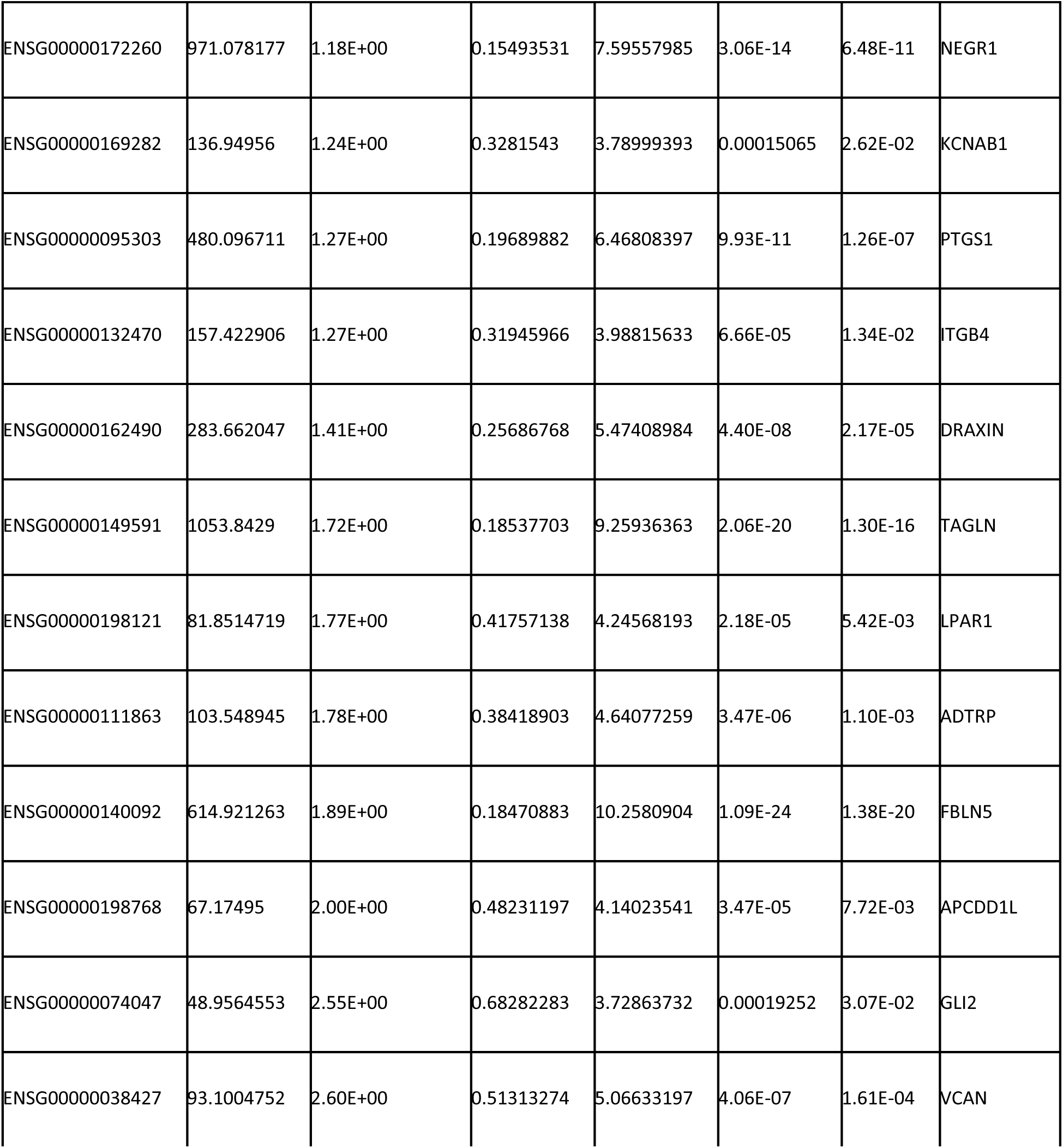
Significantly differentially regulated genes between EHBP1si and Scramble groups.

**Table S2.** List of differentially related genes containing EGF repeats or related to EGF signaling.

| Contain EGF repeats | Related to EGF signaling |
| --- | --- |
| LRRC32 | GPR39 |
| HSPG2 | EHBP1 |
| Notch2 | CXCL2 |
| EDIL3 | CXCL8 |
| LAMC2 | CTSS |
| PTGS1 | CXCL1 |
| ITGB4 | SEMA3G |
| FBLN5 | CCL2 |
| VCAN | THSD7A |
|  | CC2D1B |
|  | ACE |
|  | KDR |
|  | ADGRF5 |
|  | FABP4 |
|  | MAOA |
|  | FLRT2 |
|  | PPP1R16B |
|  | NFKBIA |
|  | EFNB2 |
|  | DPY19L1 |
|  | TFPI |
|  | BGN |
|  | GPR176 |
|  | HMGA2 |
|  | CD44 |
|  | IGFBP7 |
|  | MICAL1 |
|  | TNS3 |
|  | NDRG1 |
|  | EMP2 |
|  | SPOCD |
|  | NEGR1 |
|  | DRAXIN |
|  | LPAR1 |
|  | GLI2 |

